# Integrative modeling of seasonal influenza evolution via AI-powered antigenic cartography

**DOI:** 10.1101/2025.08.04.668423

**Authors:** Jumpei Ito, Shusuke Kawakubo, Hiroaki Unno, Adam Strange, Spyros Lytras, Kaho Okumura, Alice Lilley, Ruth Harvey, Nicola Lewis, Kei Sato

## Abstract

Seasonal influenza viruses evade host immunity through rapid antigenic evolution. Antigenicity is assessed by serological assays and typically visualized as antigenic maps, which represent antigenic differences among virus strains. However, conventional maps cannot directly infer the antigenicity of unexamined variants from their genotypes. Here, we present PLANT, a protein language model that projects influenza A/H3N2 viruses onto an antigenic map using HA protein sequences. Using PLANT-based cartography, we show that (i) H3N2 antigenic evolution accelerates during periods of disrupted global circulation, (ii) antigenic novelty accounts for a substantial portion of viral fitness advantage, and (iii) vaccine strains are often antigenically distant from circulating viruses. We further propose a PLANT-based framework for selecting vaccine strains with improved antigenic match than the WHO-recommended strains. This study provides a statistical foundation for integrated modeling of viral genotype, antigenicity, and fitness, offering quantitative insights into seasonal influenza virus evolution and supporting rational vaccine design.

## Introduction

One of the major challenges in controlling viral infectious diseases is the rapid evolution of viruses. Seasonal influenza viruses undergo continuous antigenic changes, enabling them to evade population immunity established by prior infection and vaccination ^1^. The emergence of immune-escape variants poses a significant risk of larger epidemics ^2–4^. Furthermore, to develop effective vaccines against circulating strains, it is essential to characterize their antigenicity and to select vaccine strains that closely match them ^1,5,6^. Therefore, the antigenicity of emerging seasonal influenza variants is continuously monitored through intensive efforts by multiple research groups, including the Worldwide Influenza Centre ^7^.

Differences in antigenicity between viral strains can be quantified by antigenic distance, which reflects the extent to which humoral immunity elicited by a reference strain can neutralize a target strain. This distance is measured using serological assays—such as neutralization or hemagglutination inhibition (HI) assays—by testing sera from animals infected with a reference strain against the target strain ^8^. Antigenic distance is defined as the difference in the log_2_ HI or neutralization titers between homologous and heterologous strain pairs ^2,8^. To comprehensively characterize the antigenicity of a given variant, it is informative to measure antigenic distances from multiple reference strains. However, the resulting data are high-dimensional and challenging to interpret intuitively. Antigenic cartography addresses this issue by embedding viral strains into a low-dimensional space using multidimensional scaling (MDS) ^8^. Antigenic cartography has been widely used to identify immune-escape variants, guide vaccine strain selection, and track antigenic evolution ^1,9^. For instance, vaccine strains have been updated to satisfy the quantitative criterion that their antigenic distance from circulating strains remains within two units ^8^.

Conventional antigenic cartography cannot be used to predict the antigenicity of strains that have not been experimentally characterized ^8^. If the antigenicity of newly emerged variants could be inferred from their genotypes, such as the sequences of viral proteins, it would allow for rapid identification of immune-escape variants based solely on viral genome surveillance data. Also, this capability could partially replace and reduce reliance on labor-intensive serological assays. Most importantly, such a predictive model would enable the construction of comprehensive antigenic maps that encompass all virus strains with available sequence data. When combined with phylogenetic and epidemic dynamics modeling, these maps could offer quantitative insights into the mechanisms of viral evolution and epidemic dynamics, while also supporting more effective vaccine strain selection.

Protein language models (pLMs) are a class of large language models trained not on natural language but on vast collections of protein sequences ^10^. The development of pLMs has enabled flexible genotype-to-phenotype modeling of proteins. Notably, pLMs embed protein sequences into a semantic space that captures structural, functional, and evolutionary characteristics of the proteins ^10,11^. Previous studies have shown that antigenic viral proteins, such as the hemagglutinin (HA) of influenza, are embedded in ways that reflect their antigenic similarity ^11^. In this study, we developed PLANT (<u>P</u>rotein <u>L</u>anguage Model for <u>Ant</u>igenic cartography), a model that predicts the antigenic map coordinates of seasonal influenza A/H3N2 viruses from the amino acid sequences of the HA1 domain, by leveraging the semantic space of a pLM. We further demonstrate the utility of PLANT-generated antigenic maps for elucidation of viral evolutionary patterns, identification of immune-escape variants, and improvement of vaccine strain selection.

## Results

### Training of PLANT

We first developed a novel algorithm we term pLM-based MDS to convert the semantic space of a pLM into an antigenic map (**Figure 1A**). In this framework, a pLM augmented with dimensionality reduction layers embeds the HA1 sequences of viral strains into a low-dimensional semantic space. In this space, pairwise distances between the embedded sequences—referred to as semantic distances—are then computed. The model is trained to minimize the discrepancy between these semantic distances and experimentally measured antigenic distances (termed observed antigenic distances). Through this training, the semantic space is transformed into an antigenic map that reflects antigenic relationships. This method can be regarded as an extension of MDS used in conventional cartography, augmented by the semantic space of a pLM. Importantly, unexamined virus strains can be embedded into the antigenic map constructed using pLM-based MDS, based on their HA1 sequences via the pLM.

**Figure 1.**
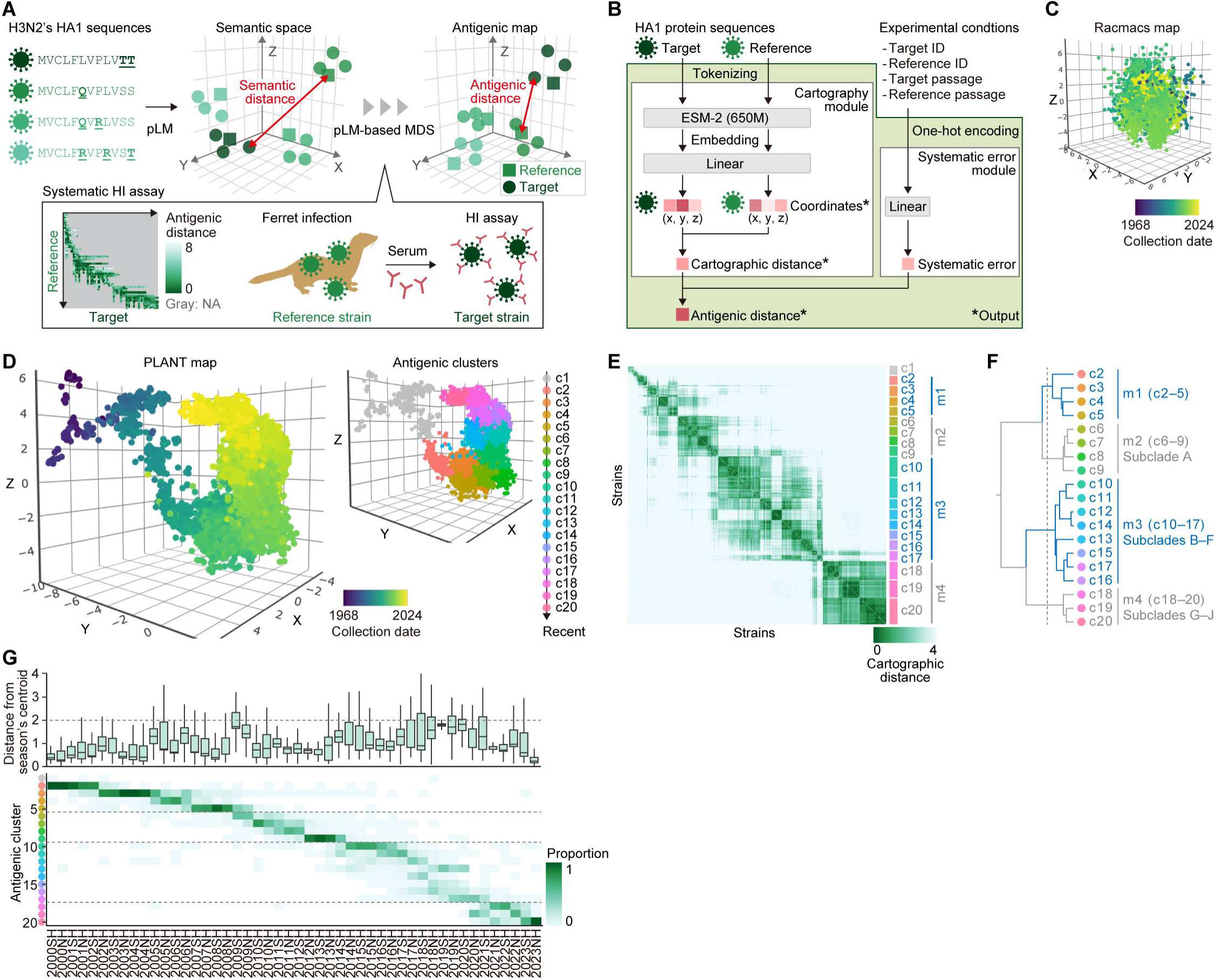
**Construction of the antigenic map by PLANT**A) Overview of PLANT training. Each virus strain is embedded in a low-dimensional semantic space derived from a pLM using its HA1 sequence. The model is trained so that the distances between target and reference strains in this space, referred to as semantic distances, reflect experimentally measured antigenic distances. This training transforms the semantic space into an antigenic map. In the bottom left, the structure of the HI assay dataset is shown. Each point represents a virus pair tested in an HI assay, colored by the observed antigenic distance. Gray indicates pairs without available measurements. Target and reference strains are ordered by collection date. B) Model architecture of PLANT. The model takes HA1 sequences of target and reference strains, along with experimental conditions, as input. It outputs coordinates in the antigenic map, the cartographic distance (Euclidean distance on the map), and antigenic distance adjusted for estimated systematic error. C) Antigenic map reconstructed using a conventional method, Racmacs (https://acorg.github.io/Racmacs/). D) Comprehensive antigenic map covering 151,784 H3N2 strains with HA1 sequences collected from 1968 to January 2024. Points are colored by collection date (left) or by 20 antigenic clusters (i.e., c1–20; right). E) Cartographic distance matrix. For each antigenic cluster, 10% of strains (or at least 20 if fewer) were randomly sampled. Strains are ordered by antigenic cluster and collection date. Antigenic metaclusters defined in (**F**) are labeled. F) Hierarchical clustering of antigenic clusters. Based on centroid coordinates, clusters were grouped into four metaclusters (i.e., m1–4) using Ward’s method with a defined cutoff (dotted line). Antigenic cluster c1, which consists primarily of pre-2000 strains, was excluded. Nextclade subclade classifications ^17^ corresponding to each metacluster are also indicated. G) Distributions of cartographic distances from seasonal centroids (top) and seasonal detection frequencies of antigenic clusters (bottom). Dotted lines indicate the 2-antigenic unit criterion (top) and metacluster boundaries (bottom). SH: Southern Hemisphere season; NH: Northern Hemisphere season.

PLANT comprises two main components: a cartography module, which constructs the antigenic map using pLM-based MDS, and a systematic error module, which corrects experimental biases arising from differences in assay conditions (**Figure 1B**). The cartography module utilizes the 650-million (M)-parameter version of ESM-2 ^12^ as its pLM. Given a pair of HA1 sequences for a reference and a target strain, along with metadata describing the experimental settings, PLANT predicts their coordinates on the antigenic map, the Euclidean distance between them (termed cartographic distance), and the systematic error–corrected distance (termed antigenic distance). If an unpaired HA1 sequence is provided, PLANT returns its predicted coordinate. Unlike the conventional method ^8^, the cartographic distance predicted by PLANT is symmetric; it remains unchanged regardless of which strain is designated as the reference.

For model training, we used HI assay data for seasonal influenza A (H3N2) viruses collected between 1968 and January 2024 by the Worldwide Influenza Centre, along with corresponding HA1 protein sequences obtained from GISAID (https://gisaid.org/). The final dataset comprised 63,118 observed antigenic distances measured between 208 reference strains and 5,271 target strains (**Figure S1A**). Because HI assays are typically conducted between temporally proximate strains, the resulting antigenic distance matrix is highly sparse, with most entries missing (**Figure 1A, bottom left; Figure S1A**) ^13^. Moreover, since this dataset was constructed by combining experimental data accumulated over several decades, it inevitably incorporates various sources of systematic error, as mentioned in previous studies ^14–16^. We randomly split the dataset along the target virus axis, ensuring that no antigenic distance measurements involving the same target strain appeared in both the training and test sets. This dataset was split without considering the temporal information of virus strains and is hereafter referred to as the “full dataset.”

### Antigenic map generated by PLANT

When the conventional antigenic cartography tool Racmacs was applied to the full dataset (prior to train–test splitting), the resulting map failed to preserve the temporal continuity of antigenic evolution (**Figure 1C**). In sharp contrast, the antigenic map generated by PLANT, trained on the full dataset (after train–test splitting), successfully captured a continuous trajectory of antigenic drift spanning the entire evolutionary timeline of human H3N2 viruses (**Figure 1D, left; Table S1**). This contrast highlights the importance of incorporating viral genetic information and correcting for systematic error in order to construct effective antigenic maps from this dataset, in line with previous reports ^14^. Details of PLANT’s predictive performance are presented in a later section. Furthermore, because PLANT can predict antigenic map coordinates even for strains lacking serological measurements, it enables the construction of a comprehensive map encompassing all 151,784 H3N2 strains with fully determined HA sequences collected between 1968 and January 2024 (**Figure 1D, left**).

To characterize the discontinuous nature of H3N2 antigenic evolution, we performed clustering based on the predicted coordinates and assigned strains to 20 antigenic clusters (**Figure 1D, right**). As expected, these clusters showed strong concordance with established H3N2 classification systems, such as the Nextclade subclade designations ^17^ (**Figure S1B; Table S1**). Since Nextclade designations are unavailable for older strains (e.g., those sampled before 2010), our downstream analyses primarily relied on the antigenic cluster classification. We next constructed a distance matrix representing pairwise cartographic distances between strains. This analysis confirmed that strains within the same antigenic cluster were tightly co-localized in the antigenic space (**Figure 1E**). Furthermore, hierarchical clustering based on cluster centroids revealed a higher-order structure, grouping 19 clusters (excluding c1, which mainly comprised pre-2000 strains) into four metaclusters: m1 (c2–5), m2 (c6–9), m3 (c10–17), and m4 (c18–20) (**Figure 1F**). Metaclusters m2, m3, and m4 corresponded to subclades A, B– F, and G–J in Nextclade, respectively (**Figure S1B**). Together, these results demonstrate that the PLANT-derived antigenic map effectively reconstructs over five decades of H3N2 antigenic evolution, capturing both gradual drift and discrete changes between major antigenic lineages.

We found that antigenic diversity varied substantially across seasons, as quantified by the cartographic distances from the centroid of epidemic strains in each season (**Figure 1G, top; Table S1**). Notably, seasons marked by the co-circulation of multiple antigenic clusters— such as the 2009 Southern Hemisphere (SH; February–August) season and the 2019 Northern Hemisphere (NH; September–January) season—exhibited pronounced increases in antigenic diversity, with median distances approaching 2 antigenic units (**Figure 1G, bottom**). These findings suggest that during such antigenically diverse seasons, selecting a single vaccine strain that is antigenically proximate to all circulating variants may be inherently difficult. In contrast, antigenic diversity declined following the emergence of rapidly expanding lineages, which swept through the population and displaced pre-existing strains.

### Prediction performance of PLANT

We first evaluated the predictive performance of PLANT trained on the full dataset, which had been trained using a randomly split dataset based on target virus strains, as described earlier.

Prediction accuracy, measured by mean absolute error (MAE) and Pearson correlation coefficient, was assessed for each influenza season from 2010SH to 2023NH (**Figure S2A**). In the analyzed dataset, HI assay results showed a mean standard deviation of 0.42 among replicates conducted under identical experimental conditions, reflecting the intrinsic noise of the assay. Nevertheless, PLANT achieved a seasonal average MAE of 0.83 and a correlation coefficient of 0.78, indicating robust predictive performance. Moreover, PLANT’s antigenic distance predictions outperformed both the cartographic distances (i.e., without systematic error correction) and the scale-adjusted semantic distances derived from the non-finetuned ESM-2 (**Figure S2A**). These results demonstrate that, under the interpolation condition, PLANT can accurately predict the antigenicity of previously unseen virus variants.

To evaluate the extrapolation performance of PLANT, we next assessed its ability to predict the antigenicity of future variants (**Figure 2**). Based on virus collection dates, the dataset was divided into “past” and “future” season datasets (**Figure 2A**). PLANT was trained exclusively on the past dataset and evaluated on the future dataset. This procedure was repeated for 19 different cutoff seasons, ranging from 2014SH to 2023SH, following the settings in Shah et al.^15^. We compared PLANT’s performance against two benchmarks: (i) a PLANT model trained on the full dataset, and (ii) AdaBoost, a state-of-the-art model for antigenicity prediction developed by Shah et al ^15^. Like PLANT, AdaBoost predicts antigenic distances from HA1 sequences and experimental metadata, but unlike PLANT, it cannot predict an antigenic map.

**Figure 2.**
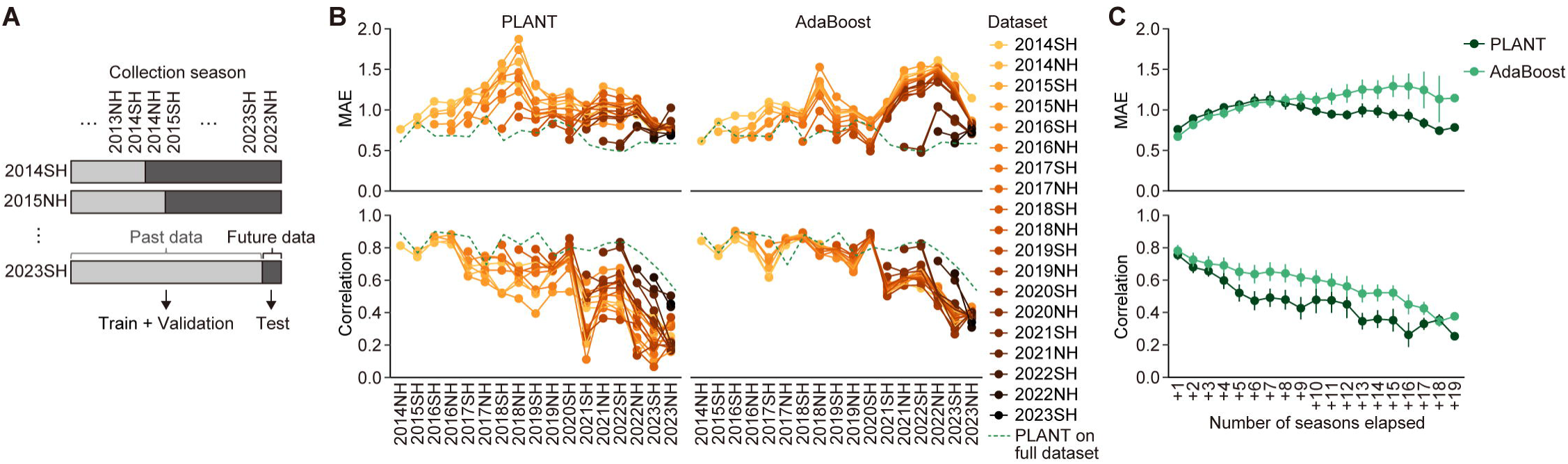
Evaluation of the extrapolation performance of PLANT. A) Dataset splitting based on virus collection date. For each cutoff (e.g., end of 2014SH), virus strains were split into past and future subsets. PLANT was trained on past data and evaluated on future data. Nineteen such datasets were prepared using cutoffs from 2014SH to 2023SH. B) Summary of prediction performance across all cutoff points. For each later season, MAE (top) and Pearson correlation (bottom) are shown for PLANT and an AdaBoost model ^15^. Results from PLANT trained on the full dataset are overlaid for comparison. Only seasons with HI assay data for ≥20 target strains are included. C) Average prediction performance stratified by the number of seasons since the cutoff. Error bars indicate standard deviations.

As expected, both PLANT and AdaBoost trained only on past data showed reduced accuracy compared to the PLANT model trained on the full dataset, and prediction accuracy generally declined with increasing temporal distance from the cutoff season (**Figures 2B, 2C, S2B, and S2C**). Comparing PLANT and AdaBoost, we found that their MAE values were similar in early post-cutoff seasons. However, after +7 seasons (approximately 3.5 years), AdaBoost’s performance declined (i.e., higher MAE) markedly, while PLANT maintained stable accuracy (**Figure 2C**). This decline was particularly evident after the 2021NH season, coinciding with the emergence of an antigenically distinct viral lineage, metacluster m4 (**Figures 2B and 1G**). These findings suggest that PLANT outperforms AdaBoost in MAE-based performance for distant future variants. For the Pearson correlation coefficient, both models performed similarly in early post-cutoff seasons (within +3 seasons). However, in later seasons, PLANT’s correlation tended to decline more than AdaBoost’s. Taken together, these results indicate that PLANT achieves extrapolation performance comparable to that of the existing state-of-the-art model, at least for several seasons into the future.

### Rate of antigenic evolution

Accurately estimating the rate of viral antigenic evolution is crucial for understanding the epidemic factors and mutations driving antigenic change, as well as for optimizing the frequency of vaccine strain updates. To investigate this rate in human H3N2 viruses, we performed an integrative analysis combining PLANT-based antigenic maps with phylogenetic modeling. Specifically, we projected the time-calibrated phylogenetic tree of H3N2 viruses onto the PLANT-based antigenic map and inferred branch-specific rates of antigenic evolution (**Figures 3A and 3B**).

**Figure 3.**
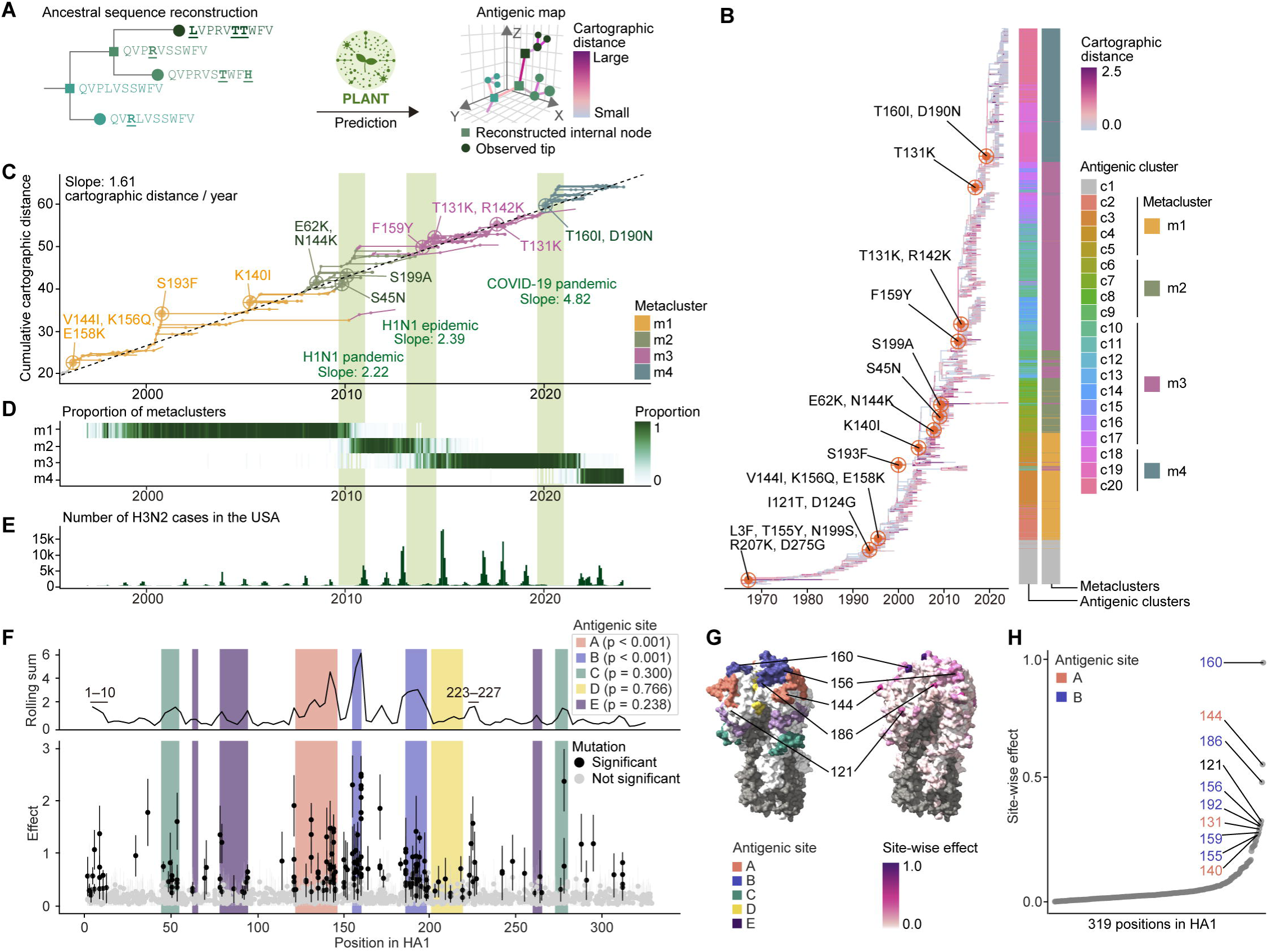
Rate of antigenic evolution and genetic drivers of antigenic change in H3N2. A) Workflow for estimating antigenic changes along phylogenetic branches. Following ancestral sequence reconstruction, PLANT (trained on the full dataset) was used to predict antigenic map coordinates for all tip and internal node sequences. Cartographic distances between parent and child nodes were then calculated for each branch. B) Time-scaled maximum-likelihood phylogenetic tree of H3N2 HA1, with branches colored according to the magnitude of antigenic change. A total of 2,748 HA1 sequences, downsampled via 99% sequence identity clustering, were included. Antigenic clusters and metaclusters are annotated at the tip nodes. Internal nodes exhibiting significant antigenic changes are labeled with their corresponding acquired substitutions. C) Antigenic evolution along the phylogenetic trunk, defined as the set of internal branches with ≥10 descendant tips. The y-axis indicates the cumulative cartographic distance along each node from the root. Antigenic cluster c1 was excluded. Branch color indicates antigenic metaclusters. Internal nodes associated with major antigenic transitions are annotated with the acquired substitutions. The periods corresponding to the 2009 H1N1 pandemic, the 2013–2014 H1N1 epidemic, and the COVID-19 pandemic are highlighted. Evolutionary rates of antigenic change for the indicated periods, estimated by linear regression, are shown. D) Epidemic dynamics of antigenic metaclusters. The proportion of each metacluster in viral genome surveillance is shown for each monthly time bin. E) H3N2 epidemic curve (monthly reported cases) based on FluNet data from the US (https://www.who.int/tools/flunet). F) Estimated antigenic effects of individual mutations using a Bayesian non-negative linear model. The lower panel shows the posterior means and 94% highest density intervals (HDIs). Mutations whose 94% HDI lower bounds exceed 0.05 are considered to have statistically significant effects and are shown in black. Colored regions represent antigenic sites A–E ^43^. The P values for each antigenic site were calculated by comparing the effect sizes of mutations within the site to those outside the site using a two-sided Mann–Whitney U-test, followed by multiple testing correction with the Benjamini–Hochberg method. The upper panel shows the rolling sum of estimated effects across sequence positions (window size = 5, step size = 3). G) HA trimer structure [PDB ID: 4FNK] ^47^, visualized by known antigenic sites (left) and by site-wise effect sizes. Site-wise effects were calculated as the sum of mutational effects at each site, scaled by a factor of 1/20. H) Ranking of amino acid positions based on site-wise effects.

To estimate the average rate of antigenic evolution in H3N2, we calculated the cumulative antigenic change along the branches comprising the trunk of the phylogeny— defined as the set of branches with ≥10 descendant tips—from the root (**Figure 3C**). We then performed a linear regression to model the cumulative cartographic distances of both external and internal nodes as a function of their sampling or inferred dates. This analysis revealed an average antigenic evolution rate of 1.61 antigenic units per year. Although this estimated slope is slightly higher than previously reported estimates (e.g., 1.2 antigenic units in Smith et al. ^8^), this discrepancy may stem from differences in the time periods analyzed, as the earlier study focused on an older dataset (i.e., 1968–2003).

Consistent with the pattern observed in **Figure 1E**, the rate of antigenic change during H3N2 evolution appears to vary over time (**Figure 3B**). By assessing antigenic change on the H3N2 phylogeny, we found that external branches tend to accumulate a greater degree of antigenic changes compared to internal branches (**Figure S3A**). This suggests that a substantial proportion of mutations leading to larger antigenic change are not transmitted to subsequent lineages, likely due to their deleterious effects ^18^. Furthermore, some of the internal branches exhibited markedly elevated antigenic changes, with 12 branches showing cartographic distances exceeding the three standard deviations over the mean (1.65 units). As expected, some of these corresponded to antigenic cluster transitions (**Figure 3B**). Notably, the rate of antigenic evolution increased substantially at the boundaries between antigenic metaclusters (**Figures 1F, 3C, and 3D**). These transitions occurred during 2009NH–2010NH (m1–m2 boundary), 2013SH–2014SH (m2–m3 boundary), and 2019NH–2020NH (m3–m4 boundary), with corresponding antigenic evolution rates reaching 2.22, 2.39, and 4.82 units per year, respectively (**Figure 3C**). Interestingly, periods of accelerated antigenic evolution coincided with major interruptions in global H3N2 circulation—specifically, the 2009 H1N1 pandemic, the 2013–2014 H1N1 epidemic, and the COVID-19 pandemic (**Figure 3E**). Among these, the most pronounced acceleration occurred during the COVID-19 pandemic, which marked the most severe interruption of global influenza circulation ^19^. These observations raise the possibility that such large-scale interruptions in continuous H3N2 circulation may facilitate or accelerate antigenic evolution (see **Discussion**).

### Mutations driving antigenic evolution

To identify amino acid substitutions responsible for antigenic changes during H3N2 evolution, we estimated the effect of individual substitutions on cartographic distance by analyzing a phylogenetic tree embedded in the antigenic map. Cartographic distance is defined as non-negative, and we assumed that the total antigenic change between a parent and its descendant strain results from the sum of non-negative contributions of individual amino acid substitutions, in accordance with Neher et al. ^2^. Based on this assumption, we implemented a Bayesian non-negative linear model in which antigenic change per branch is explained as a non-negative linear combination of individual mutation effects (**Figure S3B**). This analysis identified 195 amino acid mutations across 85 sites as having statistically significant contributions to antigenic change (**Figures 3F and S3C; Table S2**).

As expected, the majority of these mutations were located within antigenic sites known to be targeted by neutralizing antibodies (**Figures 3F–H; Table S3**). In particular, antigenic sites A and B showed significant enrichment of impactful mutations, consistent with a previous study ^20^. Furthermore, significant mutations were identified at all seven positions (145, 155, 156, 158, 159, 189, and 193) previously proposed to drive antigenic cluster transitions ^20^. Moreover, significant mutations were also observed outside these canonical antigenic sites.

For example, positions 1–10 and 223–227—regions encompassing the undefined epitopes proposed by Shih et al. ^21^—were enriched for significant mutations, supporting a potential role for these regions in antigenic evolution. Together, these results demonstrate that PLANT can effectively capture the impact of individual mutations to antigenic changes and thus serves as a powerful tool for dissecting the genetic basis of antigenic evolution.

### Relationship between antigenic novelty and fitness advantage

Using PLANT, we constructed antigenic maps solely from genomic sequence data. This enabled us to reconstruct both the epidemic dynamics and antigenic evolution of viruses from the same genome surveillance data, thereby allowing the integrated modeling of viral genotype, antigenicity, and fitness.

Using this framework, we first examined the relationship between *antigenic novelty* and *fitness advantage*. Variants (i.e., groups of strains) that are antigenically distinct from previously circulating strains can generally evade population immunity. Such variants with higher antigenic novelty are thought to exhibit higher fitness advantage enabling them to outcompete co-circulating variants ^1^. However, the quantitative relationship between antigenic novelty and fitness advantage—defined as the extent to which a variant’s fitness exceeds that of other concurrently circulating variants—remains poorly understood (**Figure 4A**). Therefore, we applied our integrative modeling approach using PLANT to address this issue.

**Figure 4.**
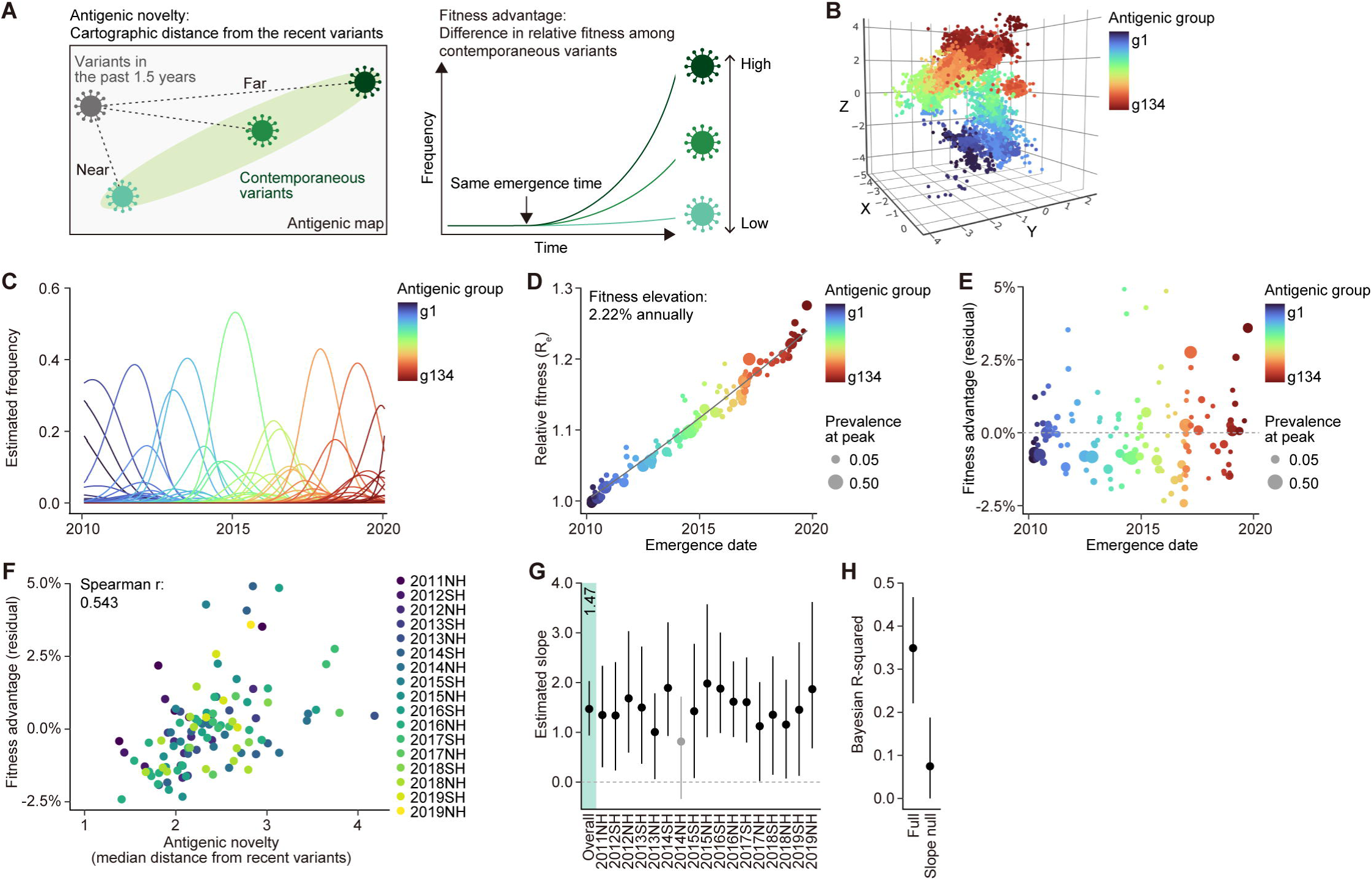
**Relationship between antigenic novelty and fitness advantage**A) Antigenic novelty and fitness advantage. Left) Antigenic novelty, using cartographic distance from recently circulating strains as a proxy. Right) Fitness advantage, using difference in relative fitness (effective reproduction number; R_e_) among contemporaneous variants as a proxy. B) Definition of antigenic groups by GMM clustering on the antigenic map. Virus strains from 2010SH to 2019NH were classified into 134 antigenic groups (g1–134). C) Epidemic dynamics of antigenic groups estimated using a Bayesian multinomial logistic model. Lines indicate posterior means. D) Emergence date and relative R_e_ of each antigenic group. Emergence date is defined as the 5th percentile of strain collection dates. Relative R_e_ values—normalized to the dominant 2010NH strain—are shown as posterior means. Point size reflects peak prevalence. The fitted exponential regression curve and the estimated average annual increase in relative R_e_ are also indicated. E) Emergence date and fitness advantage of each antigenic group. Fitness advantage was defined as the residual from the exponential regression in (**D**) and expressed as a percentage. F) Association between antigenic novelty and fitness advantage. The x-axis represents antigenic novelty, defined as the median cartographic distance from strains that circulated within the previous 1.5 years. G) Effect size of cartographic distance on fitness advantage estimated via Bayesian hierarchical modeling. Posterior means and 90% credible intervals are shown for each season and overall (highlighted in light green). Gray indicates seasons when the lower 5% of the posterior is below zero. H) Bayesian R^2^ values ^46^ for the model. Posterior means and 95% credible intervals are shown. As a control, the R^2^ value for a model that does not incorporate cartographic distance effects is also included.

The fitness of a variant can be evaluated based on its effective reproduction number (R_e_), which represents the virus’s growth potential in a population over time ^22–27^. By applying a multinomial logistic model to time-series data of variant frequencies derived from viral genome surveillance, the relative R_e_ values between variants can be estimated ^22–27^. Although this method assumes that the relative fitness of variants remains constant over time—a simplification that may not always reflect real-world dynamics—it is a widely used approach for comparing the fitness of co-circulating variants based solely on genome surveillance data ^22–27^.

We first classified H3N2 strains into *antigenic groups*—variants defined based on antigenicity—using a Gaussian mixture model (GMM) clustering on the PLANT-derived antigenic map, resulting in 134 groups (g1–134; **Figure 4B**). The analysis focused on the span between the 2010SH and 2019NH seasons, before the epidemic interruption by the COVID-19 pandemic. We then estimated their epidemic dynamics and their relative R_e_ using a Bayesian multinomial logistic model (**Figure 4C**). As expected, antigenic groups that emerged later tended to exhibit higher R_e_, reflecting temporal replacement dynamics and aligning with findings from previous studies ^25,26^ (**Figure 4D**). An exponential regression analysis showed that the R_e_ of H3N2 viruses has increased by approximately 2.22% per year.

As a proxy of antigenic novelty, the median cartographic distance from prior epidemic strains collected within the preceding 1.5 years was quantified. This time window was selected to capture strains from the most recent epidemic season in the same hemisphere. Fitness advantage was defined as the difference between a variant’s relative fitness and the expected value based on its date of emergence. Accordingly, we used the residual from the exponential regression curve shown in **Figure 4D** as a proxy for fitness advantage (**Figure 4E**).

Correlation analysis revealed a positive association between antigenic novelty and fitness advantage (Spearman’s ρ = 0.543; **Figure 4F**). Using a hierarchical Bayesian model, we then estimated the effect size both as an average across seasons and for each season individually (**Figure 4G**). The effect size was positive in all seasons except 2014NH, and an overall mean estimate was 1.47. This implies that a one antigenic unit increase in antigenic novelty corresponds to an average relative R_e_ increase of 1.47%. Combining this with the estimated antigenic evolution rate (1.61 antigenic units per year; **Figure 3C**), the annual increase in relative R_e_ attributable to antigenic change is inferred to be at 2.37% (1.47% per antigenic unit × 1.61 antigenic units per year), which closely aligns with the directly estimated 2.22% increase in **Figure 4D**. Furthermore, the Bayesian R-squared of the model was 0.35, suggesting that 35% of the measured variation in fitness advantage can be explained by antigenic novelty, together with season-specific effects (**Figure 4H**). These findings collectively suggest that antigenic change is a major driver of replacement epidemic variants in H3N2 influenza viruses.

### Automated identification of emerging immune-escape variants

We further applied the integrative framework that combines PLANT with epidemic dynamics modeling to automatically identify variants with both distinct antigenicity and high epidemic potential. This method follows the scheme described above: constructing an antigenic map using PLANT, classifying virus strains into antigenic groups, and estimating the relative R_e_ of each group. Because this approach defines variants (i.e., antigenic groups) based on antigenicity and estimates their fitness accordingly, it eliminates the need for experts to manually define variants or cross-reference fitness estimates with an antigenic map.

In this analysis, PLANT was trained on data up to the 2023NH season, and the antigenic map was constructed using viruses collected through the 2024SH season (**Figures 5A–C**). This one-season delay in PLANT’s training data reflects the typical lag in the availability of serological assay results. Using this setup, virus strains sampled between the 2022SH and 2024SH seasons were classified into 30 antigenic groups (g1–30).

**Figure 5.**
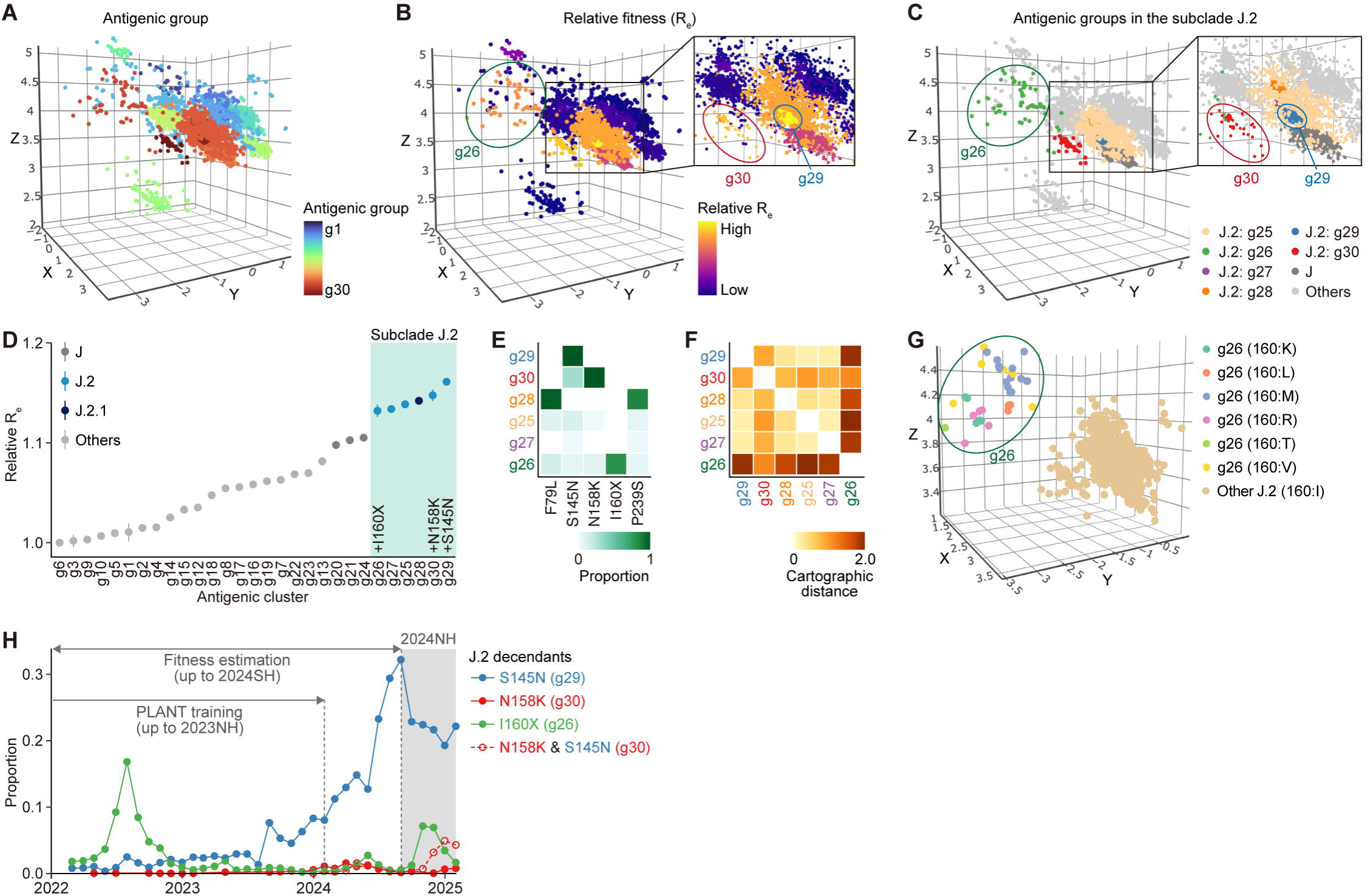
Automated identification of upcoming immune escape variants with PLANT. A) Antigenic group classification. PLANT trained on data through 2023NH was applied to viruses collected between 2022SH and 2024SH (note: 2024SH was not included in training). GMM clustering was used to define 30 antigenic groups (i.e., g1–30). B) Antigenic map colored by estimated relative R_e_ of each antigenic group. C) Visualization of antigenic groups of interest. D) Relative R_e_ values of antigenic groups. R_e_ is scaled to 1 for g6 (dominant in 2022SH). Posterior means and 95% credible intervals are shown. E) Amino acid substitution patterns for the top six antigenic groups ranked by estimated fitness. Amino acid substitutions are relative to A/Darwin/6/2021. Only substitutions with differing presence or absence patterns across the antigenic groups are displayed. F) Cartographic distance matrix among centroids of clusters in the J.2 subclade. G) Antigenic map colored by amino acid residues at position 160. Only J.2 subclade strains are shown. H) Observed epidemic dynamics of J.2 descendants carrying specific mutations, based on viral genome surveillance data up to 2024NH. Proportions were calculated in monthly bins. The period during which data were collected for PLANT training and fitness estimation is annotated.

The top six antigenic groups ranked by relative R_e_ exhibited significantly higher R_e_ values than the remaining groups (**Figure 5D**). According to the Nextclade subclade classification, all six high-fitness groups belonged to subclade J.2 (including J.2.1), which had been rapidly expanding and had become the dominant lineage as of July 2025 (https://nextstrain.org/seasonal-flu/h3n2). Notably, these J.2-related groups exhibited distinct mutational profiles, with several harboring key amino acid substitutions known to contribute to antigenic change (**Figure 5E**). For example, antigenic groups g29, g30, and g26—ranked 1st, 2nd, and 6th in fitness, respectively—contained the substitutions S145N, N158K, and I160X (X denoting any amino acid). Some strains in g30 carried S145N in addition to N158K, the hallmark mutation of this group. These mutations are revertant substitutions that occurred at known hotspots for recurrent mutations during H3N2 virus evolution ^1,20^. On the antigenic map, antigenic group g29 (rank 1) was located near other J.2 lineage viruses, whereas g30 (rank 2) and g26 (rank 6) were positioned more distantly, suggesting that they are antigenically distinct from the main J.2 lineage (**Figures 5C and 5F**). In particular, antigenic group g26 was located farthest from the other J.2 strains, consistent with the fact that g26 carries substitutions at position 160, in which substitutions showed the highest estimated impact on antigenicity (**Figure 3H**). Also, isoleucine at position 160 is a hallmark of antigenic cluster c20 and was acquired on the stem branch of this cluster (**Figure 3C**). Antigenic group g26 comprises strains bearing various amino acids at position 160, suggesting that the loss of isoleucine at this site plays a key role in antigenic change, regardless of the substituting residue (**Figures 5G**).

Together, these findings indicate that subclade J.2 encompasses multiple antigenically divergent lineages.

Finally, to assess whether these J.2 descendant strains carrying distinctive mutations actually expanded, we analyzed viral genomic surveillance data from the 2024NH season, which were used in neither PLANT training nor fitness estimation (**Figure 5H**). The J.2 lineage carrying the S145N substitution (J.2+S145N), corresponding to antigenic group g29—the variant with the highest estimated fitness—began expanding in 2023NH and accounted for >30% of circulating strains at the beginning of 2024NH. In contrast, J.2+I160X (g26) and J.2+N158K&S145N (part of g30) remained at low prevalence throughout the 2023NH and 2024SH seasons, making signs of expansion difficult to detect visually. However, both lineages subsequently expanded after 2024NH, eventually comprising approximately 5% of circulating strains at their respective peaks. The J.2+N158K lineage, which also falls within g30, remained at low prevalence by the end of 2024NH. Collectively, these findings demonstrate that the PLANT-based framework enables the early and automated identification of antigenically novel variants with the potential for epidemiological expansion.

### Evaluation of historical vaccine strains

Vaccine strain selection involves choosing the most appropriate strain—among those circulating up to a given point in time—to serve as the antigen in upcoming vaccination campaigns. For seasonal influenza, vaccine strains are updated up to twice per year in response to antigenic changes in circulating viruses ^1,6,7^. A key challenge in this process is the substantial lead time required for vaccine production and distribution, which necessitates selecting a strain that is predicted to be antigenically similar to viruses expected to circulate seven to 12 months in the future. For the Northern and Southern Hemispheres, vaccine strains are recommended each February and September, respectively, through WHO-coordinated Vaccine Composition Meetings ^1,6,7^.

To systematically evaluate the historical vaccine strains, we calculated the cartographic distance between each virus and its nearest vaccine strain used since 2001, based on the antigenic map generated by PLANT trained on the full dataset (**Figures 6A and 6B; Table S1**). This analysis revealed that 97.2% of virus strains were located within 2 antigenic units of their closest vaccine strain. While some strains exceeded this threshold, recent strains isolated after 2020 were underrepresented among them (**Figure 6B**). These results suggest that updates to the H3N2 vaccine strain have successfully tracked the major trajectory of antigenic evolution.

**Figure 6.**
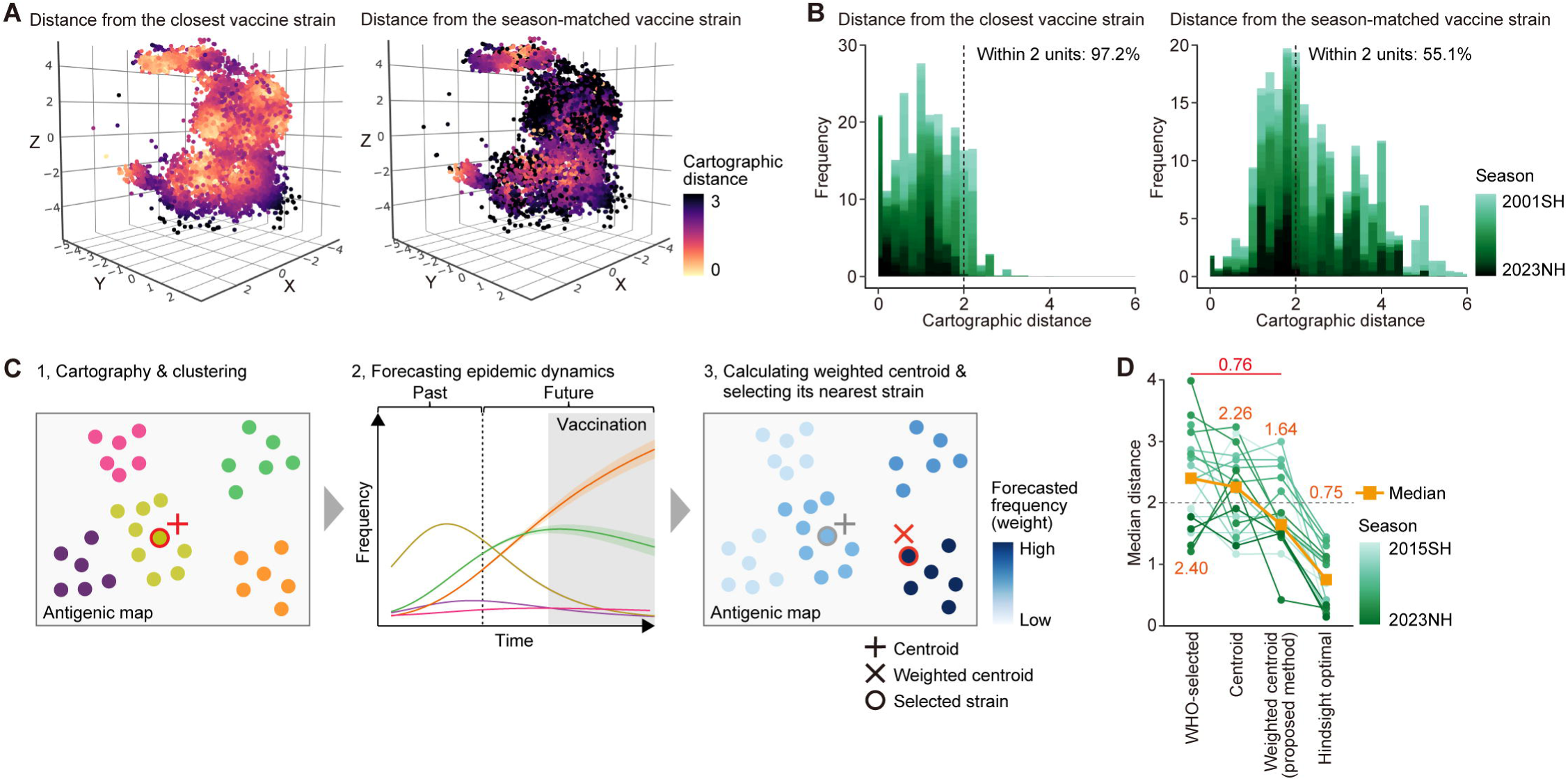
Evaluation of historical vaccine strains and vaccine strain recommendation with PLANT. A) Cartographic distance from each virus strain to the nearest historical vaccine strain (left) and to the season-matched vaccine strain (right). Analysis includes viruses from 2001SH onward. B) Histogram of cartographic distances to the nearest historical vaccine strain. The proportion within 2 antigenic units is indicated. C) Vaccine strain recommendation framework. 1) Construct an antigenic map for the last five seasons and assign viruses to antigenic groups. 2) Predict the prevalence of antigenic groups during the vaccination period using a multinomial logistic model. 3) Compute a weighted centroid and select the nearest strain as the vaccine strain. The predicted prevalence of antigenic groups during the vaccination period was used as the weight. D) Comparison of selected vaccine strains based on antigenic match, assessed by the median cartographic distance to strains circulating during the vaccination period. This panel compares (i) WHO-recommended strains, (ii) strains nearest to the centroid of the most recent season, (iii) strains selected by our proposed framework (weighted centroid), and (iv) hindsight-optimal strains. The orange line and accompanying numbers indicate the overall median across all seasons. The difference between WHO-selected strains and those chosen by our framework is highlighted in red. The dotted line denotes the 2-antigenic unit threshold.

In sharp contrast, when cartographic distances were calculated between each virus and the vaccine strain actually used in the same season, only 55.1% of viruses fell within 2 antigenic units. Furthermore, even among viruses isolated after 2020, 20.3% of strains were more than 2 antigenic units away from the corresponding vaccine strain. These findings indicate that vaccine strain updates have not consistently kept pace with the rapid antigenic evolution of H3N2, resulting in frequent mismatches at the time of vaccination.

### An optimized framework for vaccine strain selection

To address the above issue, we developed a new vaccine strain selection scheme that integrates PLANT with epidemic dynamics modeling (**Figure 6C**). In this framework, PLANT is trained annually using genomic and HI assay data available as of January for the Northern Hemisphere and August for the Southern Hemisphere. Based on the trained model and sequence data available by those dates, we construct an antigenic map, classify viruses into antigenic groups, and use a multinomial logistic model to forecast the epidemic dynamics of each group during the vaccination period. The forecasted prevalence of each antigenic group during the vaccination period is then computed, and these values are used to assign weights to virus strains on the antigenic map. A weighted centroid is computed, and the strain closest to this centroid is selected as the candidate vaccine strain.

We evaluated the performance of our vaccine selection scheme through retrospective simulations spanning the 2015SH to 2023NH seasons. As a proxy for antigenic match, we computed the median cartographic distance between the selected vaccine strain and the viruses that circulated during the vaccination period, based on the antigenic map by PLANT trained on the full dataset. These distances were compared across four types of strains: (i) the actual WHO-recommended strains, (ii) the strains nearest to the centroid of the most recent season, (iii) the strains selected by our proposed framework, and (iv) hindsight-optimal strains—defined as those closest to future epidemic strains, selected using future information (**Figure 6D**).

Among the strategies compared, the WHO-recommended strains satisfied the 2-antigenic unit criterion in 8 out of 18 seasons, with a median antigenic distance of 2.40 units (**Figures 6D and S4A**). The nearest-centroid strains exhibited a similar median distance. In contrast, our weighted centroid approach met the threshold in 12 seasons and achieved a substantially lower median distance of 1.64 units. The lowest distance (0.32 units) was observed in the 2022NH season, due to the accurate prediction of the expansion of the newly emerged antigenic cluster c20 (**Figure S4B**). Although the performance of our method does not yet match the hindsight-optimal benchmark of 0.75 units, it represents an average reduction of 0.76 antigenic units compared to the WHO-recommended strains. Given that a one-unit antigenic distance corresponds to a two-fold change in neutralization activity ^8^, this improvement translates to an estimated 1.69-fold increase in vaccine-induced neutralization. Taken together, these findings demonstrate that integrating our automated scheme provides a promising strategy for improving the accuracy and effectiveness of H3N2 vaccine strain selection.

## Discussion

We here present PLANT, a protein language model that projects influenza A/H3N2 viruses onto an antigenic map using only HA1 sequences. Unlike conventional cartography methods, PLANT enables antigenic mapping for all sequenced strains. This capability allows for the direct integration of antigenic cartography with genomic surveillance through phylogenetic and epidemic modeling. Using this unified framework, we demonstrate that: (i) H3N2 antigenic evolution accelerates during periods of disrupted global circulation (**Figure 3**); (ii) antigenic change is a major driver of variant replacement in H3N2 (**Figure 4**); (iii) emerging immune-escape variants with high epidemic potential can be identified automatically at an early stage (**Figure 5**); and (iv) vaccine strain selection can be made more predictive and effective (**Figure 6**). Together, these results highlight PLANT as a comprehensive platform for jointly modeling viral genotype, antigenicity, and fitness—advancing both our understanding of antigenic evolution and our capacity to control viral infectious diseases.

Although previous studies have developed numerous statistical and machine learning models to predict antigenic distances between pairs of virus strains, these models cannot directly generate antigenic maps ^2,3,14–16,28,29^. As an *ad hoc* solution, some studies have used the predicted distances to construct antigenic maps using conventional cartography methods based on MDS ^2,15,28,29^. This approach, however, requires reconstructing the entire map whenever new strains are added, leading to inconsistencies in the coordinate system across updates. PLANT overcomes these limitations by directly predicting antigenic map coordinates from HA1 sequences, enabling real-time monitoring of viral antigenic evolution in conjunction with routine viral genome surveillance. To realize this capability, we developed a novel learning algorithm, pLM-based MDS, which trains a pLM directly from pairwise antigenic distances and sequence data—without relying on pre-built antigenic maps generated by conventional cartography methods. Beyond antigenic mapping, pLM-based MDS provides a general framework for transforming the semantic space of a pLM into a supervised phenotypic space. This approach opens new avenues for flexible genotype–phenotype modeling with pLMs.

The association between antigenic change and fitness advantage among variants has long been recognized in seasonal influenza viruses, particularly H3N2 ^1^. However, previous analyses have largely relied on (i) a limited set of variants with available antigenic measurements, (ii) genetic distance as a proxy for antigenic distance, or (iii) season-level analyses based on average values of antigenic change and fitness advantage ^2–4,23^. In this study, we conducted the first comprehensive assessment of the relationship between antigenicity and fitness advantage at the variant level (**Figure 4**). We estimated that a one-unit increase in antigenic novelty corresponds to a 1.47% increase in relative fitness, with a Bayesian R-squared value of 0.35. Although this indicates a substantial association, the effect size—as represented by the slope—was modest. This reflects the inherently limited annual fitness increase observed in H3N2, estimated at approximately 2.22% per year (**Figure 4D**).

Such limited fitness gains lead to slower replacement of epidemic lineages over time, as previously discussed^1,18,30–32^. Importantly, this result suggests that immune-escape variants are unlikely to trigger exceptionally large-scale epidemic waves in H3N2. For instance, to achieve a 20% increase in fitness—commonly observed every few months in SARS-CoV-2 between 2020 and 2024 ^25,26^—a variant would require approximately 13.6 antigenic units of change, equivalent to 8.4 years of antigenic evolution in H3N2. The small effect size further supports the view that while the recurrent emergence of immune-escape variants influences the magnitude of seasonal epidemics, it is not the primary driver of their occurrence ^1–4^. These findings demonstrate that PLANT offers a quantitative understanding of the relationship between viral antigenicity and fitness advantage.

Global circulation of H3N2 has been intermittently disrupted by major respiratory epidemics, including the 2009 H1N1 and COVID-19 pandemics, as well as the 2013–2014 H1N1 epidemic ^4,19,33^. Our analysis suggests that the rate of antigenic evolution accelerated during these periods of reduced circulation (**Figures 3C–E**). Previous studies have proposed that H3N2 evolution is accelerated by population bottlenecks that usually arise during seasonal transitions, when the dominant epidemic population shifts (e.g., from the Northern to the Southern Hemisphere) ^1^. Our findings support this hypothesis and further extend it to more extreme scenarios involving global interruptions of viral circulation. The importance of bottleneck effects in H3N2 evolution likely stems from the relatively modest fitness advantages conferred by antigenic changes (**Figure 4G**). That is, because even immune-escape variants require prolonged periods to reach fixation in large populations, rapid antigenic evolution likely depends on the increased stochasticity introduced by bottlenecks ^30–32^. Also, previous studies proposed that the fixation of advantageous immune escape mutations in the evolution of H3N2 is slowed down by the accumulation of sublethal deleterious mutations, increasing fitness variance in the virus population ^18,34^. Taking this into consideration, our findings are consistent with a scenario where population bottlenecks, resulting from the reduced circulation periods, decrease fitness variance and, in turn, enable more efficient fixation of antigenic change mutations. Future studies should aim to elucidate the evolutionary dynamics and specific mechanisms underlying these patterns.

The antigenic map constructed by PLANT enables a systematic assessment of antigenic similarity between vaccine strains and circulating viruses (**Figure 6**). This property of our model can be used not only to retrospectively evaluate historical vaccine choices but also to benchmark alternative selection strategies. Indeed, we demonstrated that a simple PLANT-based framework can recommend vaccine candidates that are antigenically closer to circulating strains than those recommended by the WHO. While logistic regression is widely used to forecast the dynamics of emerging variants ^23^, our framework seamlessly integrates this approach with antigenic map predictions by PLANT, enabling the automated and efficient selection of vaccine strains. While real-world vaccine selection must also consider practical constraints—such as strain availability and growth efficiency in egg-based production ^5^—our scheme offers a principled statistical framework for optimizing strain selection. In the future, integrating our scheme with more accurate, machine learning–based epidemic forecasting methods could further improve vaccine strain selection.

This study has several limitations. First, the extrapolation performance of PLANT for entirely novel future variants remains limited and does not clearly surpass that of the state-of-the-art AdaBoost model ^15^ (**Figure 2**). Nevertheless, the key strength of PLANT lies in its ability to construct antigenic maps and perform downstream quantitative analyses under interpolative conditions. Moreover, because PLANT can be retrained within a few hours on a single GPU, regularly updating the model with the latest data reduces reliance on extrapolation and ensures adaptability to recent antigenic trends. Second, PLANT defines cartographic distances symmetrically, meaning that a given genotype is mapped to the same coordinates regardless of whether it is treated as a reference or a target strain. While this differs from the original asymmetric implementation ^8^, we adopted the symmetric formulation based on preliminary experiments that showed improved extrapolation performance. Third, although we conducted retrospective validation using temporally split datasets, further prospective validation with newly emerging viral sequences will be necessary to rigorously assess PLANT’s practical utility in real-time surveillance and vaccine strain selection.

Despite these limitations, our findings highlight the utility of PLANT not only in advancing our understanding of antigenic evolution but also in improving the control of viral infectious diseases. Importantly, the PLANT framework is not limited to seasonal influenza; it is broadly applicable to other viral pathogens for which serological data are available, such as SARS-CoV-2, dengue virus, and highly pathogenic avian influenza, and is extendable to future pandemic threats. By jointly modeling viral genotype, antigenicity, and fitness, PLANT provides a powerful foundation for understanding, forecasting, and ultimately mitigating viral epidemics.

## Methods

### Dataset preparation

We retrieved all available HA sequences of human H3N2 influenza viruses and their associated metadata from GISAID (https://gisaid.org/) as of January 28, 2025. The translated amino acid sequences of the HA1 domain were extracted and aligned using Nextclade v3.14.0 ^17^, with A/Wisconsin/67/2005-egg (GenBank accession number: CY163680.1) as the reference. Clade and subclade classifications were also assigned by Nextclade. We excluded sequences from non-human hosts, sequences lacking collection date information, and those containing undetermined amino acids in the HA1 domain. Also, only HA1 sequences with exactly 329 amino acids were retained.

In addition, we collected HI assay data and related experimental metadata from the annual and interim reports of the Worldwide Influenza Centre (https://www.crick.ac.uk/research/platforms-and-facilities/worldwide-influenza-centre/annual-and-interim-reports). The raw data up to the 2021NH season were derived from Harvey et al. ^16^, and those through the 2024NH season were generously provided by the Worldwide Influenza Centre.

HI assays were stratified into experimental batches based on the date of experiment and the passage type of the reference strain, and observed antigenic distances were computed within each batch. HI titers (*T_AB_*) were measured for combinations of target strain A and reference strain B, forming a matrix of HI measurements with not assigned (NA) values, indicating strain pairs for which HI assays were not performed (**Figure S1A**). The observed antigenic distance between strains A and B was calculated using the following formula as described in previous studies ^2,8^: *D_AB_* = log_2_(*T_AB_*) - log_2_(*T_BB_*). The observed antigenic distance represents the number of two-fold dilution steps by which the titer for the heterologous strain A differs from that of the homologous strain B, both measured against sera from ferrets infected with the reference strain B. Observed antigenic distance is thus an integer value.

In some cases, the observed antigenic distance was negative, indicating a higher titer for the heterologous strain pair than for the homologous strain pair—likely due to experimental noise. To avoid potential adverse effects of such noise on model training, measurements with values below –1 were excluded, while those equal to –1 were set to 0.

Each HI measurement was matched to its corresponding HA1 protein sequence via strain names. After filtering, the final dataset comprised 63,118 observed antigenic distances, measured between 208 reference strains and 5,271 target strains (**Figure S1A**). Observed antigenic distances ranged from 0 to 8, with some values right-censored (e.g., >6).

Before training PLANT, the observed antigenic distances were scaled to a range between 0 and 1. After inference, the predictions were rescaled to the original scale by multiplying by a scaling factor of 8.

For the unsupervised training of PLANT, we constructed a dataset of HA protein sequences spanning diverse HA subtypes of influenza A viruses from both human and animal hosts. HA protein sequences were retrieved from the NCBI Influenza Virus Database (https://www.ncbi.nlm.nih.gov/genomes/FLU/Database/nph-select.cgi). All unique type A sequences at the amino acid level were downloaded, resulting in a total of 33,657 sequences (retrieved on April 11, 2024). Metadata including serotype and collection date were also obtained for all sequences. To remove redundancy, the dataset was clustered using MMSeqs2 ^35^ with a minimum sequence identity threshold of 99% (--cov-mode 0), and a single representative sequence was retained from each cluster. Sequences containing ambiguous characters (X or J) and those annotated as mixed serotypes were further excluded, resulting in a final HA-all training dataset comprising 9,890 sequences. The HA1 domain was extracted from each sequence and used for downstream analysis.

### Overview of PLANT

PLANT is a model that embeds each virus strain into a Euclidean space based solely on its HA1 protein sequence. In this space—hereafter referred to as the antigenic map—virus strains with identical HA1 sequences are assigned the same coordinates, regardless of whether they serve as reference or target strains. The distance between strains, referred to as the cartographic distance, is defined as the Euclidean distance between their embedded coordinates in the antigenic map. Cartographic distance is a symmetric, non-negative value that depends solely on the HA1 sequence.

In contrast, antigenic distances measured by serological assays are influenced not only by true antigenic differences but also by variation in experimental conditions and measurement error. Also, true antigenic differences are potentially affected by genotype differences of viral components other than HA1, such as the HA2 domain or neuraminidase (NA). In this study, we assume that observed antigenic distances consist of the sum of three components: a cartographic distance derived from the HA1 sequence; a systematic error associated with experimental conditions, including antigenic differences arising from non-HA1 viral components; and a random error component.

During training, PLANT learns to match the observed antigenic distance with the sum of a semantic distance (i.e., the distance between sequence embeddings in the dimension-reduced semantic space computed by a pLM) and the estimated systematic error. Through this process, the semantic distance is transformed into a cartographic distance, and the semantic space is converted into an antigenic map space.

In addition to supervised learning based on pLM-based MDS, PLANT incorporates a semi-supervised learning framework by utilizing unlabeled HA1 sequences (i.e., sequences without associated serological data). By imposing structural constraints on the antigenic map through unsupervised losses, this framework enhances the generalization ability of the model (see **Section Unsupervised learning loss**).

For inference, PLANT takes as input a pair of HA1 protein sequences (reference and target strains) along with a set of categorical variables representing experimental conditions. It computes antigenic map coordinates for both sequences, calculates the cartographic distance, and adds the estimated systematic error to produce the predicted antigenic distance. If only an unpaired sequence (e.g., a target strain) is provided, the model outputs its coordinate in the antigenic map.

### Technical discussion on pLM-based MDS

A recent study ^36^ reported a model that, like PLANT, predicts the antigenic map coordinates of virus strains from viral sequences. However, this model adopts a learning approach that targets the coordinates constructed by conventional cartography methods as prediction outputs. As shown in **Figure 1C**, conventional cartography fails to accurately reconstruct the overall trajectory of H3N2 antigenic evolution, revealing a structural limitation of this approach. In contrast, our study introduces a new learning framework, pLM-based MDS, which embeds virus strains into a low-dimensional space based on pairwise antigenic distances, without relying on a preconstructed antigenic map. This method transforms the semantic space of a pLM into a phenotypic space under supervision using antigenic distances between sequence pairs. pLM-based MDS can be extended beyond antigenic mapping to construct diverse phenotypic spaces, offering a new direction for flexible genotype–phenotype modeling using pLMs.

### Architecture of PLANT

PLANT comprises two modules: (i) a cartography module that computes antigenic map coordinates from HA1 sequences and outputs the corresponding cartographic distance, and (ii) a systematic error module that estimates the component of the observed antigenic distance not attributable to the HA1 genotype, based on experimental metadata.

The cartography module consists of a pre-trained pLM, ESM-2 (650M parameters) ^12^, which takes tokenized HA1 sequences as input, followed by three fully connected layers. The fully connected layers project the per-sequence embedding (CLS token embedding) from EMS-2 into a 3-dimensional coordinate. The cartographic distance is computed as the Euclidean distance between the coordinates of the reference and target strains.

The systematic error module receives four categorical variables as one-hot encoded inputs: the IDs of the target and reference strains, and the passage types (egg, cell, and unknown) of the target and reference viruses. These metadata are used to statistically account for sources of antigenic variation not attributable to the HA1 genotype, including differences in HA2 or NA, variation in virus stock lots, infectivity, hemagglutination properties, and host animal effects. The selection of metadata follows a prior work ^15^.

The module systematic error consists of two subnetworks: (i) a three-layer fully connected subnetwork that takes the reference strain ID and both passage types as input, and (ii) a single-layer subnetwork that takes only the target strain ID. The outputs of the two subnetworks are summed to yield the systematic error. This modular design improves prediction stability for target strains not included in the training data (i.e., those with missing IDs).

### Supervised learning loss

To train PLANT on observed antigenic distances between reference–target strain pairs, we designed a custom loss function that addresses three key challenges: (i) right-censoring of observed values, (ii) imbalance in the distribution of observed antigenic distances, and (iii) the need to minimize not only the discrepancy between predicted antigenic and observed distances but also that between cartographic and observed distances.

With respect to (i), in HI assays, antigenic distance measurements are subject to an upper bound, and some observations are right-censored ^8^. Accordingly, we used a conditional loss function: for uncensored data, the loss was the squared error between predicted and observed values; for censored data, the squared error was applied only if the prediction fell below the observed value, and set to zero otherwise. This design penalizes underestimation while allowing overestimation for censored observations.

With respect to (ii), because large antigenic distances are rare, the distribution of values is heavily imbalanced. To mitigate this, we divided the range 0–8 into nine bins, assigned each bin a weight proportional to the inverse of its frequency (capped at 5), and used these weights to compute a weighted squared error across all training samples.

With respect to (iii), in PLANT, the primary objective is not to perfectly predict observed antigenic distances but to construct an antigenic map that faithfully reflects underlying antigenic differences. Therefore, we included an additional regularization term that minimizes the squared error between the cartographic distance and the observed antigenic distance. This term was weighted by 0.05 and added to the total loss.

### Unsupervised learning loss

To improve the generalizability of PLANT by leveraging information on the diversity and sequence characteristics of HA1 proteins, we incorporated unlabeled HA1 sequences—those lacking serological assay data—into model training. For this purpose, PLANT introduces two regularization losses: the semantic loss and the local-global loss, which impose semantic and geometric constraints on the structure of the antigenic map space. As unlabeled sequences, we used HA1 sequences from human H3N2 viruses, as well as those from other HA subtypes isolated from both human and animal hosts (**see Preparation of dataset for unsupervised learning section**).

The semantic loss penalizes discrepancies between semantic distances derived from the frozen ESM-2 (650M) model and the cartographic distances predicted by PLANT. To compute semantic distances, we first obtained per-sequence embeddings by applying both mean and max pooling to the final-layer outputs of ESM-2. For each mini-batch—including reference, target, and unlabeled strains—we then calculated pairwise Euclidean distances between embeddings. These distances were rescaled using a learnable scaling parameter and compared to PLANT-predicted cartographic distances via mean squared error. This loss encourages the preservation of the original semantic structure encoded by ESM-2 and helps mitigate catastrophic forgetting during fine-tuning.

The local-global loss is designed to promote both local smoothness and global separation within the antigenic map. For each sample in a mini-batch, the model minimizes the distance to its three nearest neighbors, encouraging antigenically similar sequences to be placed close together. Conversely, a penalty is applied to pairs of non-neighboring samples that lie within a margin of 0.125 (equivalent to 1 antigenic unit), promoting clearer separation between distinct antigenic clusters.

The training dataset consists of both labeled reference–target pairs (with observed antigenic distances) and unlabeled HA1 sequences. For labeled data, both supervised and unsupervised losses were applied, whereas for unlabeled data, only the unsupervised losses were used. The final training loss is a weighted sum of all loss components, with weights of 0.2 for the semantic loss and 0.01 for the local-global loss.

### Training with the full dataset

For model training using the full dataset, we split the data into training, validation, and test sets at a ratio of 0.8: 0.1: 0.1. The split was performed at the *target strain* level to ensure that all samples derived from a given target strain were assigned exclusively to either the training, validation, or test set. Stratified sampling was applied based on the collection year of the target strains to preserve temporal diversity.

The HI dataset included multiple biological replicates—that is, samples with identical virus strain, virus passage type, reference strain, and reference passage type, but different experimental dates. To reduce redundancy, we applied a downsampling strategy that randomly selected one sample from each replicate group during every training epoch.

As noted above, the training dataset included both labeled data (reference–target pairs with observed antigenic distances) and unlabeled data (individual HA1 sequences). These were randomly sampled and jointly included in each mini-batch.

Training was conducted using a custom implementation of the Hugging Face *Transformers* Trainer, with the AdamW optimizer. The parameters of ESM-2 were not frozen; full fine-tuning was performed. The learning rate was set to 1.0 × 10^-4^ and updated with a linear scheduler with a warm-up ratio of 0.1. A batch size of 16 and a total of 20,000 training steps were used. Weight decay was set to 0.005 for the fully connected layers in the cartography module and 0.01 for the other components. A dropout rate of 0.05 was applied to all fully connected layers in the cartography and systematic error modules. Validation loss was evaluated every 1,000 steps, and the model with the lowest validation loss was selected as the final model.

Training was conducted using a single NVIDIA H100 GPU (CUDA v12.8) on a high-performance computing cluster.

### Construction of a comprehensive antigenic map covering all sequenced H3N2 strains

We constructed a comprehensive antigenic map encompassing all H3N2 virus strains collected between 1968 and January 2024. Using the PLANT model trained on the full dataset, we predicted antigenic map coordinates for each virus based on its HA1 sequence. Visualization of the resulting map revealed a strong correlation between antigenic coordinates and collection dates. However, a small number of strains showed collection dates that were notably inconsistent with their predicted antigenic positions, suggesting potential metadata errors.

To detect and exclude such outliers, we implemented an outlier detection procedure (**Figure S1C**). Specifically, we used a generalized additive model (GAM) implemented in the R package mgcv v1.9 to predict collection dates from antigenic coordinates. Strains whose documented collection dates deviated by more than 20 years from the GAM-predicted values were classified as outliers and excluded from downstream analyses. After filtering, the final antigenic map included 151,784 virus strains.

### Definition of antigenic clusters

We defined 20 antigenic clusters using a two-stage Gaussian mixture model (GMM) clustering approach. In the first stage, virus strains were clustered based on their antigenic coordinates using the GMM implementation in the R package mclust v6.1.1. The number and shape of clusters were automatically determined based on the Bayesian Information Criterion (BIC), with the number of clusters explored in the range of 200 to 1000 (in increments of 100).

In the second stage, the centroid coordinates of the first-stage clusters were subjected to another round of GMM clustering, with the number of clusters explored in the range of 20 to 40 (in increments of 5). The resulting clusters from this second stage were defined as antigenic clusters and used in subsequent analyses.

### Evaluation of PLANT’s predictive performance for future strains

To assess PLANT’s ability to predict the antigenicity of novel, future-emerging variants, we split the dataset based on virus collection dates, constructing a training dataset composed of past variants and a test dataset comprising future variants.

Given that seasonal influenza vaccine strain selection meetings are typically held in February and September, we defined influenza seasons as follows: the Southern Hemisphere (SH) season spans February to August, and the Northern Hemisphere (NH) season spans September to January of the following year. For each of the 19 seasons from 2014SH to 2023SH, virus strains collected up to the cutoff season were included in the train-validation dataset, while those collected after the cutoff were assigned to the test dataset. For instance, when the cutoff is 2014SH, the train-validation dataset excludes any data with a virus or reference strain collection date on or after September 1, 2014. This temporal split was applied consistently to both labeled data (i.e., reference–target pairs with observed antigenic distances) and unlabeled data (i.e., HA1 sequences without serological measurements).

Within the train-validation dataset, we further divided data into training and validation subsets at a 0.9: 0.1 ratio. The split was stratified by the collection year of the target strains and grouped at the target strain level, following the same procedure as used in full dataset training.

### Training of the AdaBoost model

To provide a comparative benchmark for evaluating PLANT’s predictive performance, we trained the AdaBoost-based model for H3N2 antigenicity prediction proposed by Shah et al. ^15^ using the same neutralization assay dataset as PLANT. Like PLANT, the AdaBoost model takes as input the HA1 protein sequences of the reference and target strains, along with categorical variables representing experimental conditions, to predict the antigenic distance under those conditions. However, unlike PLANT, it neither predicts antigenic map coordinates nor enables the construction of an antigenic map.

Model training involved a two-step hyperparameter optimization process consisting of an initial broad grid search followed by a refined Bayesian search. Both searches were conducted using Scikit-learn’s GridSearchCV and BayesSearchCV modules on the 2014SH season dataset. In the grid search, we explored combinations of four hyperparameters: the number of estimators (n_estimators) was set to 50, 100, or 200; the learning rate (learning_rate) was set to 0.001, 0.01, 0.1, or 1.0; the maximum tree depth (estimator max_depth) was set to 500, 1000, or 2000; and the maximum number of features considered at each split (estimator max_features) was set to “auto”, “sqrt”, or “log2”. The grid search used five-fold cross-validation with negative mean absolute error as the scoring metric and was parallelized using 14 jobs. The best-performing configuration from the grid search was: n_estimators = 200, learning_rate = 0.001, estimator max_depth = 500, and estimator max_features = “sqrt”.

The Bayesian optimization then fine-tuned the search space based on these results. It sampled n_estimators as an integer between 150 and 600, learning_rate as a real number between 0.0005 and 0.005, and estimator max_depth as an integer between 100 and 750, while fixing estimator max_features to “sqrt”. This second-stage search ran for 50 iterations with five-fold cross-validation and used the same scoring metric. It was parallelized with 5 jobs. The best parameters identified were: n_estimators = 361, learning_rate = 0.0005, estimator max_depth = 728, and estimator max_features = “sqrt”.

All computations were performed on Ubuntu Linux 20.04 using Python 3.10.13. The code used for AdaBoost hyperparameter tuning and model training is available in the PLANT code repository.

### Identifying phylogenetic branches with elevated antigenic change

We analyzed HA1 protein sequences of human H3N2 influenza viruses collected between 1968 and 2024, as described in the Dataset preparation section. To reduce redundancy while preserving genetic diversity, the sequences were clustered using CD-HIT v4.8.1 ^37^ with a 99% identity threshold, yielding 2,748 representative sequences. A maximum-likelihood phylogenetic tree was then reconstructed using IQ-TREE2 v2.4.0 ^38^ with the LG+F+G4 amino acid substitution model. This tree was subsequently rooted and time-calibrated using TreeTime v0.11.4 ^39^, which also performed ancestral sequence reconstruction for all internal nodes under default settings.

Antigenic map coordinates were predicted for both terminal and internal HA1 sequences using the PLANT model trained on the full dataset. For each phylogenetic branch, we computed the cartographic distance between its parent and child nodes. Branches with significantly elevated antigenic change were identified as those whose cartographic distance exceeded the mean across all internal branches by more than three standard deviations.

To compare antigenic change between internal and external branches while minimizing sampling bias, we repeated the entire analysis pipeline—from tree construction to antigenic change inference—using all 24,166 unique HA1 sequences (i.e., without applying 99% clustering).

To estimate the average rate of antigenic evolution in H3N2, we calculated the cumulative antigenic change along the phylogenetic trunk, defined as the set of branches with ≥10 descendant tips, starting from 1997. A linear regression was then performed to model cumulative cartographic distance of both internal and external nodes as a function of their sampling or inferred dates.

Clusters and metaclusters were assigned to internal nodes based on their predicted antigenic map coordinates, using the same GMM described earlier. Monthly proportions of each metacluster were calculated. U.S. H3N2 influenza case numbers were obtained from sentinel surveillance data available via FluNet (https://www.who.int/tools/flunet).

All phylogenetic tree processing was performed using ete3 v3.1.3 ^40^, and visualizations were created with ggtree v3.14.0 ^41^.

### Estimating the effects of individual mutations on antigenic change

To identify mutations contributing to the antigenic evolution of H3N2 viruses, we developed a Bayesian non-negative linear model. Cartographic distance is inherently non-negative and defined as zero when the HA1 protein sequences of two strains are identical. Based on this, we assumed that mutations either have no effect or increase this distance. To reflect this biological constraint, the model imposes non-negativity on all regression coefficients (i.e., the effect sizes of individual mutations).

In this model, amino acid substitutions acquired on each branch are encoded as one-hot vectors and used as explanatory variables, while the response variable is the cartographic distance between the parent and child nodes as predicted by PLANT. The effect size of each mutation is assigned a prior distribution defined by a non-negative half-t distribution with zero mean and a scale parameter σ, ensuring that the posterior distribution remains non-negative. Additionally, because strains with identical HA1 sequences should have zero antigenic distance, the model does not include an intercept term. This model does not account for interactions between mutations and assumes, like standard linear regression, that mutation effects are additive.

To evaluate model validity, we compared the estimated mutation effect sizes from the Bayesian non-negative linear model with those obtained from a standard (unconstrained) linear regression (**Figure S3B**). The two estimates were broadly consistent, especially for mutations with larger effects, and the non-negative model yielded coefficients that were all zero or greater, as expected.

Parameter estimation was carried out using cmdstan v2.36.0 and cmdstanpy v1.2.5, employing Markov Chain Monte Carlo (MCMC) sampling via the No-U-Turn Sampler (NUTS) implemented in Stan. Four independent MCMC chains with 3,000 steps were run. The warm-up period was set to 1,000 steps. Before using the MCMC outputs in downstream analyses, we verified convergence criteria for all parameters: R-hat ≤ 1.05, and both bulk and tail effective sample sizes ≥ 200.

To evaluate antigenic effects at the site level rather than for individual mutations, we calculated the site-wise effect for each HA1 position by aggregating the inferred antigenic effects of all mutations occurring at the same site. Specifically, we summed the posterior mean effect of each mutation at a given position and scaled the total by a factor of 1/20, corresponding to the number of possible amino acid types.

The HA trimer structure [PDB ID: 4FNK] was visualized using ChimeraX v1.8 ^42^.

Antigenic sites were annotated based on amino acid positions defined in a previous study ^43^, and the computed site-wise antigenic effects were mapped onto the structure for spatial interpretation.

### Integrating PLANT-based antigenic cartography with epidemic modeling

Antigenic groups were defined based on the positions of virus strains in the antigenic map. To remove redundancy, we first selected a single representative from each set of viruses sharing an identical HA1 protein sequence. Clustering was then performed using a GMM implemented in the R package mclust v6.1.1, with the number of clusters automatically determined according to the BIC. For the fitness advantage analysis (**Figure 4**), we evaluated cluster numbers ranging from 100 to 500 in increments of 100. In contrast, for immune-escape variant detection (**Figure 5**) and vaccine strain selection (**Figure 6**), we used a range of 10 to 50 in increments of 10. These differences reflect variations in the number of influenza seasons analyzed—18 seasons in **Figure 4** and five seasons in **Figures 5 and 6**. The resulting antigenic group classification was applied to all virus strains, including those with redundant sequences, so that every strain was assigned an antigenic group for downstream analyses.

In the vaccine strain selection analysis (**Figure 6**), we merged antigenic groups that were excessively close in the antigenic map to achieve robust, rather than overly sensitive, groupings. For this purpose, we used the Mahalanobis distance, which accounts for correlations among dimensions and differences in scale, providing a reliable measure of multivariate distance ^44^. Specifically, for each pair of antigenic groups, we calculated the Mahalanobis distance between their centroids. If this distance was less than or equal to 1.0, the two groups were merged into a single antigenic group. This threshold of 1.0 corresponds to approximately one standard deviation in multivariate space.

For all downstream analyses, we retained only antigenic groups containing at least 20 virus strains. In the vaccine strain selection analysis (**Figure 6**), we further limited the set to those with a prevalence of at least 0.5% across the entire dataset. The emergence date of each antigenic group was defined as the 5th percentile of the collection dates of its constituent strains. Group IDs were assigned in ascending order of these emergence dates. Importantly, group IDs were dynamically assigned in each analysis and do not represent fixed or shared labels across analyses.

To estimate the epidemic dynamics and associated parameters of each antigenic group, we applied a Bayesian multinomial logistic regression model to weekly detection frequency data. This analysis was performed without country-or region-level stratification, using all available virus strains in aggregate.

Parameter estimation was carried out using cmdstan v2.34.1 and cmdstanr v2.34.1, employing MCMC sampling via NUTS implemented in Stan. Four independent MCMC chains were run for each analysis: 6,000 steps for **Figure 4** and 2,000 steps for **Figures 5 and 6**. The warm-up period was set to 500 steps in all cases. Before using the MCMC outputs in downstream analyses, we verified convergence criteria for all parameters: R-hat ≤ 1.05, and both bulk and tail effective sample sizes ≥ 200.

The relative R_e_ of the antigenic group g, 𝑟_*g*_, was calculated according to the slope parameter β_*g*_ of the antigenic group as 𝑟_*g*_ = exp(γβ_*g*_), where γ is the average viral generation time (3.0 days) ^45^. In the vaccine strain selection analysis (**Figure 6**), epidemic trajectories for each antigenic group were projected up to two seasons ahead.

### Definition of fitness advantage and its association with antigenic novelty

To estimate the fitness advantage of each antigenic group, we first classified virus strains from 2010SH to 2019NH into 134 antigenic groups using the method described above and estimated their emergence dates and relative R_e_. To capture the temporal trend in relative R_e_, we fitted an exponential regression model to the relationship between the emergence date and relative R_e_, thereby estimating the annual rate of increase in relative R_e_. The exponential regression was implemented as a linear model in which the natural logarithm of relative R_e_ served as the response variable and the emergence date as the explanatory variable. The residuals from this regression model quantify how much a group’s relative R_e_ deviates from the expected value based on its emergence date. Therefore, these residuals reflect the extent to which a given antigenic group exhibits a higher (or lower) fitness relative to contemporaneously emerging antigenic groups. This means that this residual can be interpreted as a proxy for fitness advantage.

As a proxy of antigenic novelty, the median cartographic distance from prior epidemic strains collected within the preceding 1.5 years was quantified. This time window was selected to capture strains from the most recent epidemic season in the same hemisphere.

To quantitatively assess the relationship between fitness advantage and cartographic distance, we constructed a hierarchical Bayesian linear model. In this model, we introduced season-specific random effects for both the intercept and the slope of cartographic distance, allowing us to statistically model seasonal variation. Specifically, the slope for each season was assumed to follow a normal distribution with a mean of zero and a standard deviation of σ, thereby enabling the estimation of inter-seasonal variability in a unified framework. To evaluate the explanatory power of the model, we computed the Bayesian R-squared, as defined by Gelman et al ^46^.

### Framework for automated identification of immune-escape variants

Following the procedure described above, we classified virus strains that emerged during the 2022SH and 2024SH seasons into 30 antigenic groups based on their coordinates in the antigenic map. For each group, we estimated the relative R_e_. To identify mutations potentially associated with immune escape, we used Nextclade to detect amino acid substitutions in each strain relative to the reference strain A/Darwin/6/2021 instead of A/Wisconsin/67/2005-egg.

To conduct a retrospective analysis based on virus genome surveillance data up to the 2024NH season, we downloaded HA genome sequences and associated metadata for viruses collected during this period from GISAID on June 30, 2025.

### Evaluation of historical vaccine strains and development of an automated framework for vaccine strain selection

Information on historical vaccine strains was obtained from GISAID (https://gisaid.org/resources/human-influenza-vaccine-composition/) for strains recommended from the 2010NH season onward, and from the WHO website (https://www.who.int/teams/global-influenza-programme/vaccines/who-recommendations/recommendations-for-influenza-vaccine-composition-archive) for strains from 2001SH to 2010SH. This study focused on WHO-recommended strains for egg-based inactivated influenza vaccines, which represent the most widely used modality for seasonal influenza vaccination. These WHO-recommended strains are annotated in **Table S1**.

For the labeled data used to train PLANT (i.e., observed antigenic distances derived from HI assays), we partitioned the data based on the date the HI assay was performed, rather than the collection date of the virus. For example, to train the model for predicting the vaccine strain for the 2014NH season, we included only HI assay data generated before August 31, 2013. In contrast, unlabeled HA1 sequence data were split based on the collection date of the virus.

For each cutoff date, an antigenic map was constructed using a PLANT model trained exclusively on data available up to that time point. As input, we used HA1 sequence data from viruses collected within the five preceding seasons.

Vaccine strain selection was conducted using the epidemic projection framework described earlier. Specifically, we projected the prevalence of each antigenic group up to two seasons into the future from the cutoff date and estimated the posterior mean and 50% credible interval of group prevalence during the vaccination period.

Each virus strain was then assigned a weight according to the projected prevalence (posterior mean) of its corresponding antigenic group. Using these weights, we calculated the weighted median centroid in the antigenic map and selected the virus strain closest to this centroid as the candidate vaccine strain.

To evaluate the antigenic match between the candidate vaccine strain and the circulating viruses during the vaccination period, we used the cartographic distances output by a PLANT model trained on the full dataset.

Additionally, we defined a “hindsight-optimal strain” as the virus strain that existed prior to the cutoff date and exhibited the smallest median cartographic distance to the circulating strains during the vaccination period. This allowed us to benchmark our automated selection method against the theoretical optimal choice under hindsight conditions.

## Code availability

The computational code used in this study is available at the GitHub repository (https://github.com/TheSatoLab/PLANT).

## Data availability

HI assay data and associated experimental metadata used in this study are available in the annual and interim reports of the Worldwide Influenza Centre (https://www.crick.ac.uk/research/platforms-and-facilities/worldwide-influenza-centre/annual-and-interim-reports). Genomic sequences of H3N2 viruses analyzed in this study are available from the GISAID database (https://gisaid.org/). The GISAID acknowledgment table is provided in the GitHub repository (https://github.com/TheSatoLab/PLANT).

## Acknowledgments

We gratefully acknowledge all data contributors, i.e. the Authors and their Originating laboratories responsible for obtaining the specimens, and their Submitting laboratories for generating the genetic sequence and metadata and sharing via the GISAID Initiative, on which this research is based. This study was carried out using the TSUBAME4.0 supercomputer at Institute of Science Tokyo.

We gratefully acknowledge the valuable discussions and constructive feedback provided by Dr. Junna Kawasaki (Chiba University, Japan), Mr. Takuya Hemmi (Kyoto University, Japan), Mr. Keno Strotjohann (Swiss Tropical and Public Health Institute, Switzerland), Mr. Kent Mori (Kyushu University, Japan), Mr. Moonjong Kang (The University of Tokyo, Japan), Dr. June Sese (Humanome Lab), Dr. Keietsu Abe (Tohoku University), and Dr. Ryuichi Koga (National Institute of Advanced Industrial Science and Technology).

This study was supported in part by JST PRESTO (JPMJPR22R1, to Jumpei Ito); JSPS KAKENHI Grant-in-Aid for Scientific Research B (JP25K00116, to Jumpei Ito); AMED SCARDA Japan Initiative for World-leading Vaccine Research and Development Centers “UTOPIA” (JP223fa627001, to Jumpei Ito, Kei Sato, Shusuke Kawakubo, Spyros Lytras); SHIONOGI Infectious Disease Research Promotion Foundation (to Jumpei Ito); JSPS KAKENHI Fund for the Promotion of Joint International Research (International Leading Research) (JP23K20041, to Kei Sato); AMED ASPIRE Program (25jf0126002, to Kei Sato); AMED SCARDA Program on R&D of new generation vaccine including new modality application (253fa727002, to Kei Sato); AMED Research Program on Emerging and Re-emerging Infectious Diseases (24fk0108907, 25fk0108690, to Kei Sato); AMED Japan Program for Infectious Diseases Research and Infrastructure (Collaborative Research via Overseas Research Centers) (25wm0225041, to Kei Sato); JSPS KAKENHI Grant-in-Aid for Scientific Research A (JP24H00607, to Kei Sato); the Platform Project for Supporting Drug Discovery and Life Science Research (Basis for Supporting Innovative Drug Discovery and Life Science Research (BINDS)) from AMED (JP24ama121012, supporting numbers S02820001 and S02820002, to Kei Sato), AIST KAKUSEI project (FY2024 and FY2025 to Shusuke Kawakubo).

## Author Contributions

Project design and management: Jumpei Ito Development of PLANT: Jumpei Ito Improvement of PLANT: Jumpei Ito, Adam Strange, and Shusuke Kawakubo AdaBoost-based analysis: Adam Strange PLANT-integrated phylogenetic analyses: Shusuke Kawakubo Development of a Bayesian non-negative linear model: Hiroaki Unno Immune escape variant identification analysis: Jumpei Ito and Hiroaki Unno Other analyses: Jumpei Ito Dataset preparation: Shusuke Kawakubo, Hiroaki Unno, Adam Strange, Spyros Lytras, Kaho Okumura, and Jumpei Ito Serological data provision: Alice Lilley, Ruth Harvey, and Nicola Lewis Initial draft writing: Jumpei Ito and Shusuke Kawakubo Improving the initial draft: Jumpei Ito, Spyros Lytras, and Kei Sato Figure preparation: Jumpei Ito, Shusuke Kawakubo, and Kaho Okumura Scientific illustration: Jumpei Ito, Kaho Okumura Naming of PLANT and logo design: Shusuke Kawakubo Project administration: Kei Sato All authors reviewed and approved the final manuscript.

## Supplemental Tables

**Table S1. Information on virus strains used in this study.**

The coordinates (X, Y, Z) represent the antigenic map positions predicted by PLANT trained on the full dataset. Clade and Subclade were assigned using Nextclade. The columns “is WHO vaccine strain”, “Closest vaccine strain”, “Distance from closest vaccine strain”, “Season matched vaccine strain”, and “Distance from season matched vaccine strain” are assigned only to virus strains collected from 2001SH onward.

**Table S2 Summary of predicted antigenic effects for individual mutations.**

The positions correspond to amino acid residues in the HA1 protein. The column ‘isSignificant’ indicates statistical significance, with 1 representing true and 0 representing false.

**Table S3 Summary of predicted antigenic effects aggregated by position across mutation types.**

The predicted antigenic effect of each mutation type was aggregated at each site using the sum values. The column ‘Site-wise effect’ represents the sum of effects divided by 20, corresponding to the total number of possible amino acid types at each site.

## Supplemental Figures

**Figure S1. Description of datasets (related to Figure 1)**

A) Structure of the HI assay dataset. Each point represents a virus pair tested in an HI assay, colored by observed antigenic distance. Gray color indicates virus pairs without observed antigenic distance. Target and reference strains are ordered by collection date.

B) Proportions of Nextclade subclades within each antigenic cluster. Note that subclade assignments were unavailable (i.e., NA) for most strains in clusters c1–8. Also, corresponding antigenic metaclusters are indicated.

C) Detection of virus strains with outlier collection dates. Collection dates were predicted using a generalized additive model (GAM) based on the antigenic map coordinates of each strain. Strains exhibiting a discrepancy of more than 20 years between their documented and predicted collection dates were removed as outliers. The left and right panels show the data before and after filtering, respectively.

**Figure S2. Prediction performance of PLANT (related to Figure 2)**

A) Interpolation performance of PLANT. Prediction accuracy was evaluated using random splits along the virus strain axis. Mean absolute error (MAE) and Pearson correlation are shown for antigenic distance, cartographic distance, and scaled semantic distance from the un-finetuned ESM-2, across individual seasons. Median values across seasons are shown on the right side of the plots. Only seasons with HI assay data for ≥20 target strains are included.

B and C) Evaluation of the extrapolation performance of PLANT in each epidemic season. Unlike Figure 2B, results are shown separately for 19 datasets with distinct cutoffs. For each later season, MAE (**B**) and Pearson correlation (**C**) are shown for PLANT, an AdaBoost model ^15^, and PLANT trained on the full dataset.

**Figure S3. Rate of antigenic evolution and genetic drivers of antigenic change in H3N2 (related to Figure 3)**

A) Comparison of cartographic distances between external and internal branches in the phylogenetic tree. For minimizing sampling bias, we reconstructed the phylogenetic tree of H3N2 using all 24,166 unique HA1 sequences (i.e., without applying 99% clustering).

B) Comparison of estimated substitution effects between the Bayesian non-negative linear model and the unconstrained linear model. Posterior means (dots) and 94% HDIs (lines) are shown.

C) Ranking plot of estimated substitution effects. Top 40 mutations are annotated with text labels. Mutations that are located in antigenic sites are labelled with color according to the inset color key. Significant and non-significant mutations are shown as pink and gray, respectively.

**Figure S4. Vaccine strain recommendation with PLANT (related to Figure 6)**

A) Comparison of vaccine strains selected by four different methods. Violin plots show the distribution of cartographic distances to epidemic strains during the vaccination period, with dots indicating median values.

A) Projected epidemic dynamics of antigenic groups. Results are shown for the 2022NH vaccination season. Posterior mean (line) and 50% credible interval (ribbon) are indicated. The plot includes the top five antigenic groups based on forecasted epidemic frequency during the vaccination period. For each antigenic group, the most frequent antigenic cluster and Nextclade subclade among its constituent strains are displayed alongside the group name.

