## Supplementary figures and images for "Integrative modeling of seasonal influenza evolution via AI-powered antigenic cartography"

### Figure S1

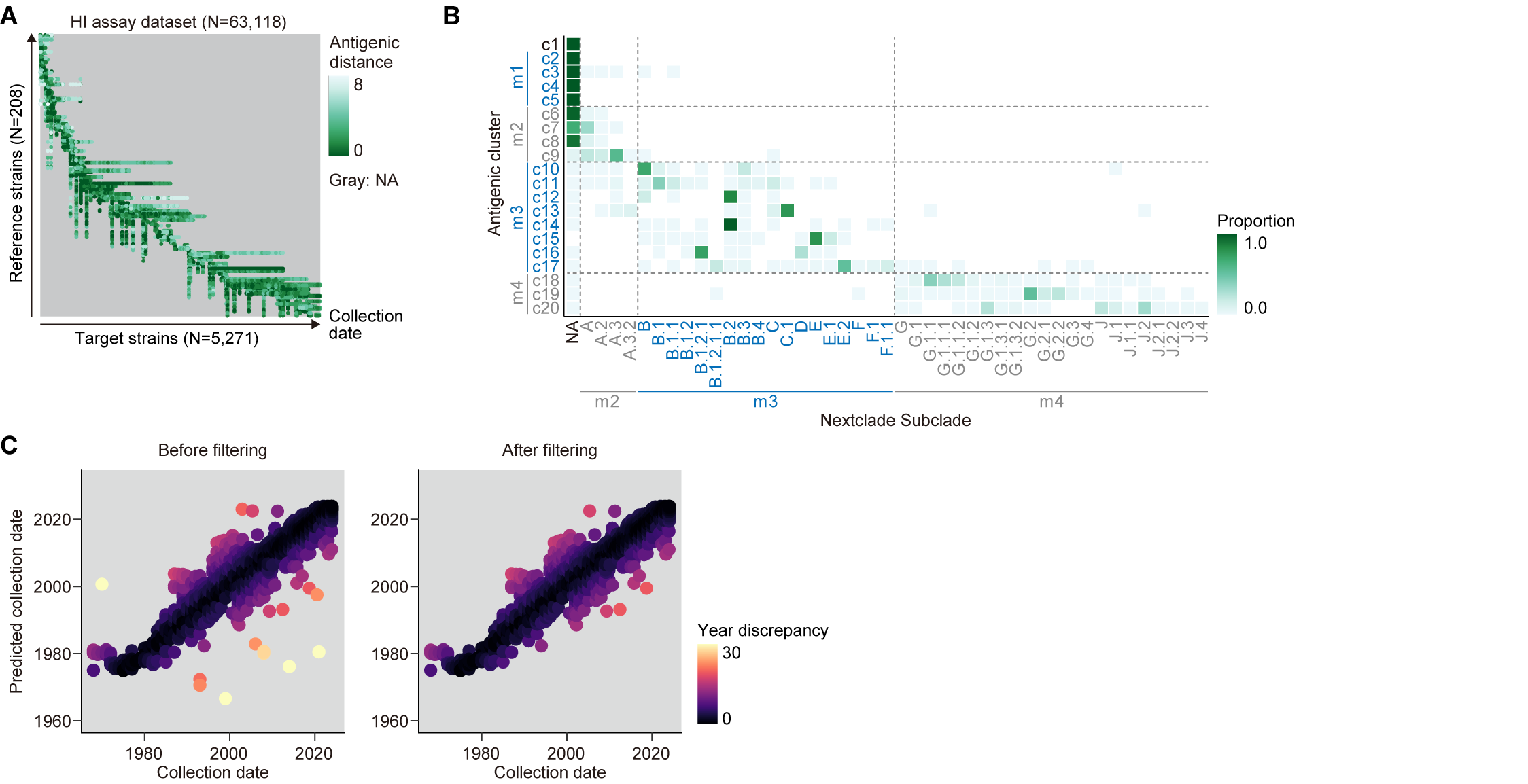

### Figure S2

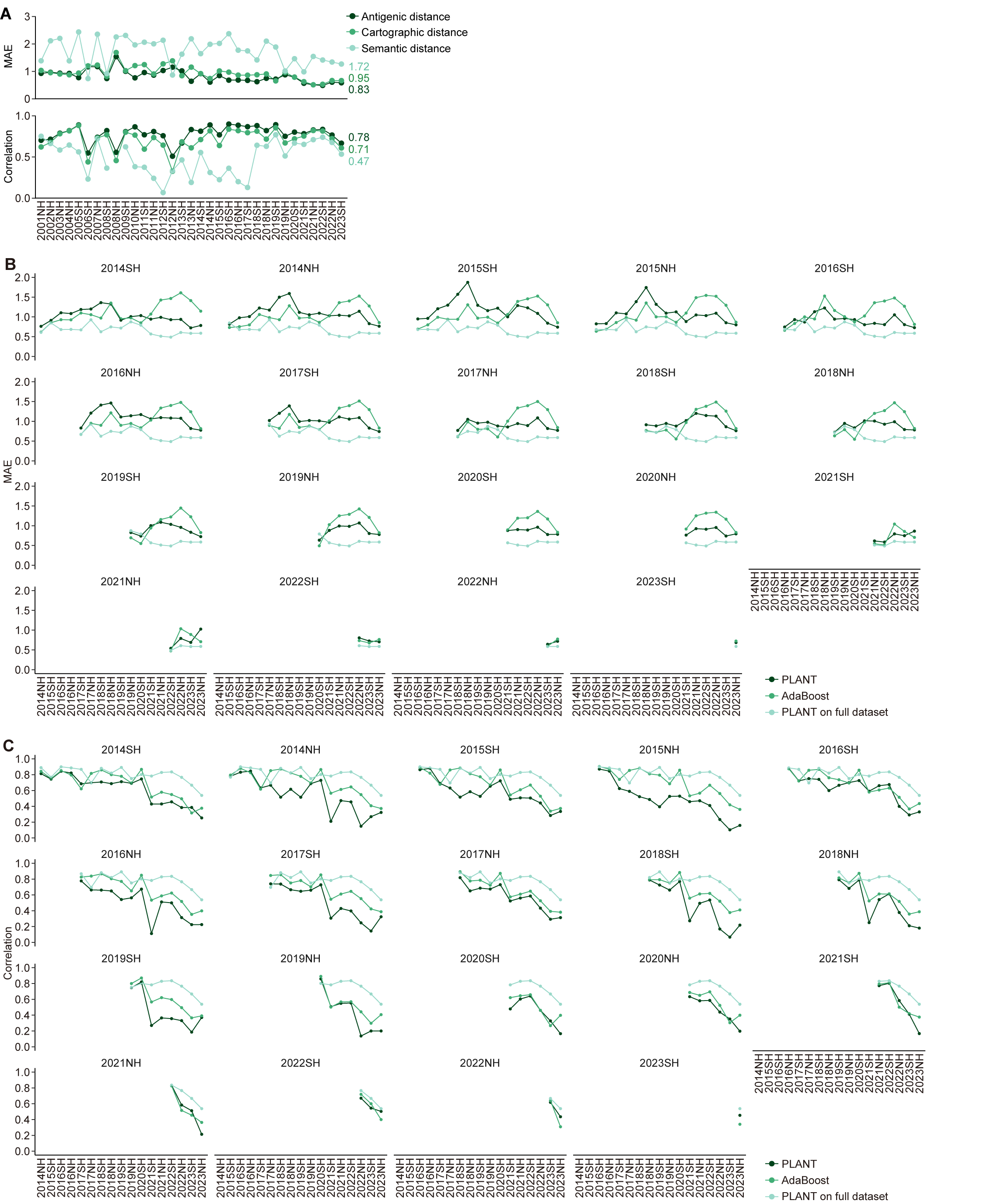

### Figure S3

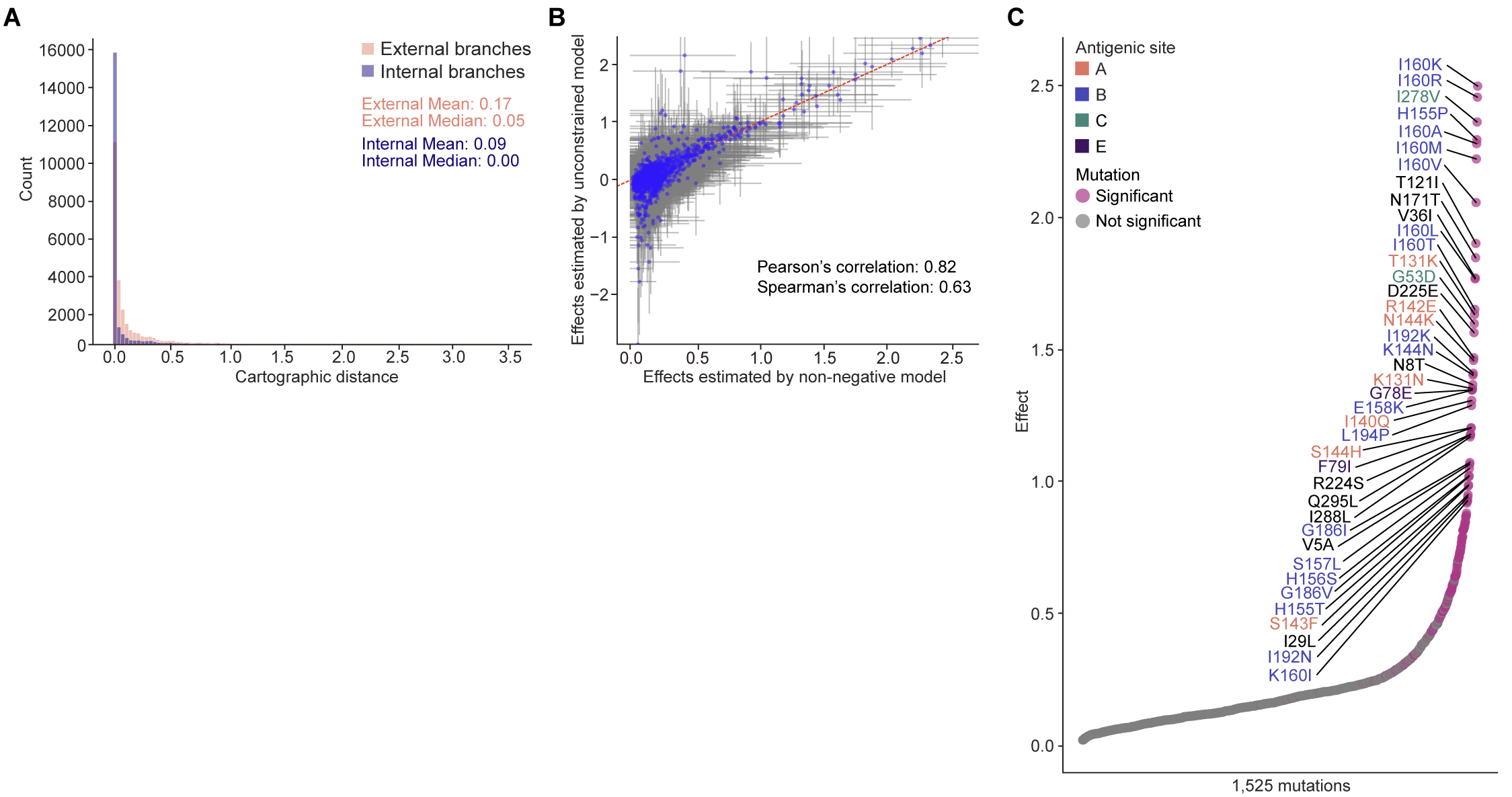

### Figure S4

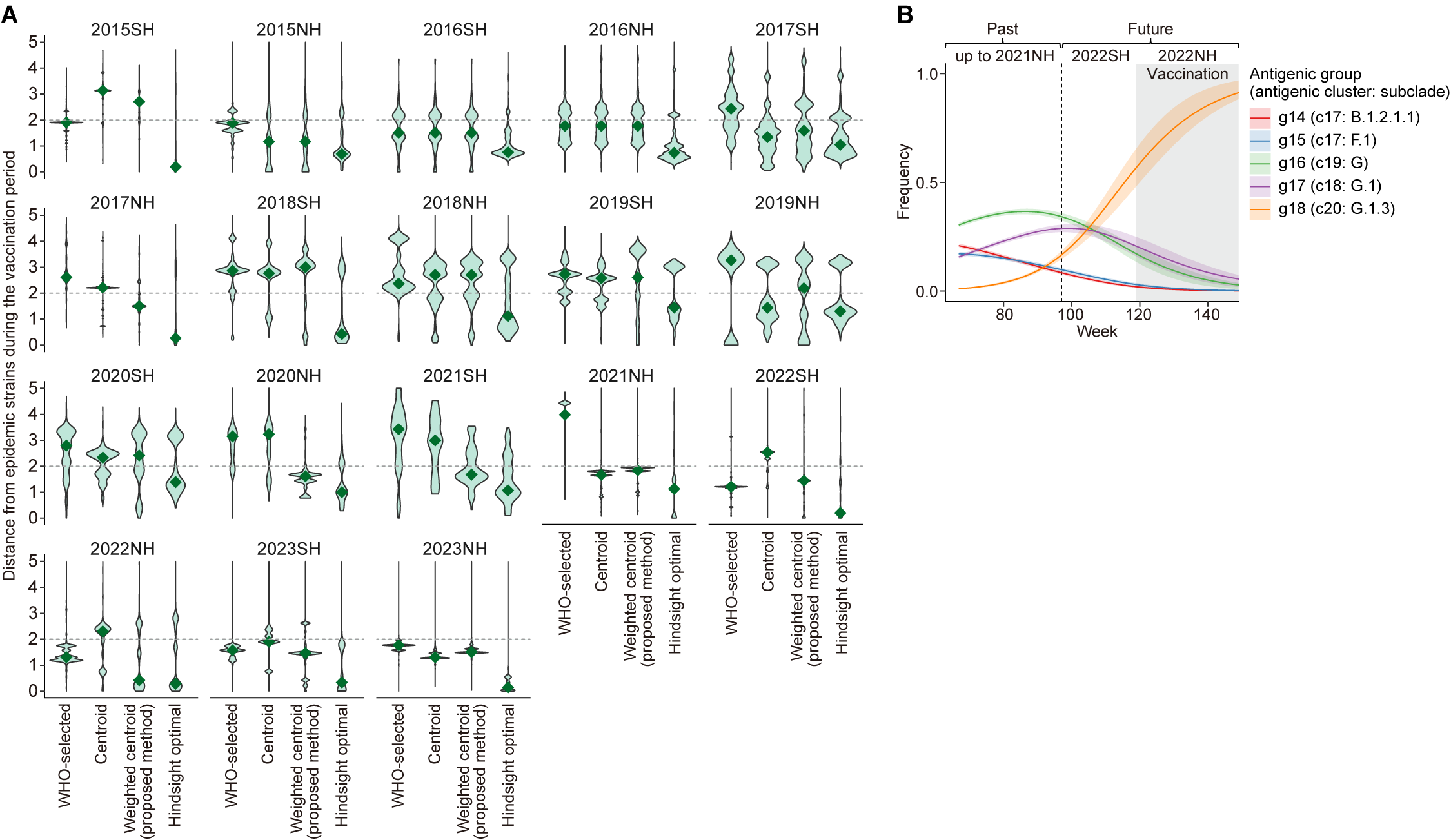
